# Balancing Efficiency and Compensation: Age-related Theta and Alpha Dynamics in High-load Working Memory

**DOI:** 10.1101/2025.11.26.690640

**Authors:** Esteban León-Correa, Philipp Ruhnau, Alex Balani, Adam Qureshi, Dorothy Tse, Stergios Makris

**Affiliations:** Edge Hill University; University of Central Lancashire; Universita degli studi eCampus

**Keywords:** Ageing, working memory, EEG, theta, alpha, oscillations, cognitive load

## Abstract

Working memory (WM) declines with age, yet older adults often preserve performance through compensatory mechanisms. Oscillatory dynamics in the theta and alpha bands are central to these adaptations, but their distinct roles across maintenance, manipulation, and recall remain unclear. We examined behavioural and neural differences in WM using electroencephalography (EEG) during verbal (Letter Span) and visuospatial (Corsi Block) tasks under retention and manipulation conditions. Behaviourally, older adults preserved performance in the Letter Span but showed deficits in the Corsi Test, suggesting domain-specific declines, while higher task demands reduced accuracy across both groups. At the neural level, younger adults exhibited stronger mid-frontal and temporal theta activity, supporting more efficient recruitment of WM processes and stronger performance associations. Older adults showed attenuated theta power, indicative of neural saturation, and appeared to compensate through alpha-mediated mechanisms during the delay period (maintenance / manipulation): reduced alpha suppression in the Letter Span facilitated rehearsal and updating, whereas increased upper alpha in the Corsi Test protected stored visuospatial information. These findings suggest theta oscillations underpin core WM functions, whereas alpha oscillations flexibly support compensatory strategies in ageing.

## INTRODUCTION

Working Memory (WM) refers to the capacity to temporarily store and manipulate information in support of goal-directed behaviour (Baddeley, 1998, 2003). It is critical for daily functioning, decision-making, and problem-solving, and is strongly associated with general fluid intelligence (Baddeley, 2012; Conway et al., 2003; Engle et al., 1999). WM is commonly divided into three phases: encoding, in which sensory input is processed and transformed into a suitable format for storage; maintenance, the active retention of information; and retrieval, the reactivation of stored content into conscious awareness before it fades (Jaffe & Constantinidis, 2021; Kim, 2019).

WM is particularly vulnerable to age-related decline and neurodegeneration (Grady, 2012; Salthouse, 2011; Ziaei et al., 2017). Older adults generally perform worse than younger adults, especially in inhibiting irrelevant stimuli and coping with interference during encoding and maintenance (Jarjat et al., 2019; Jost et al., 2011; Loaiza & Souza, 2019). Neuroimaging studies further reveal structural and functional changes that necessitate reliance on compensatory neural mechanisms to sustain performance (Cabeza et al., 2018; Rieck et al., 2017; Sala-Llonch et al., 2015). The Compensation-Related Utilisation of Neural Circuits Hypothesis (CRUNCH) and the Hemispheric Asymmetry Reduction in Older Adults (HAROLD) models propose that older adults recruit additional resources under high cognitive load but reach capacity limits earlier than younger adults (Cabeza et al., 2018; Park & Festini, 2017; Reuter-Lorenz & Cappell, 2008).

Although encoding has been extensively studied, maintenance and retrieval—where compensatory mechanisms may be most evident—remain less well understood. Comparing retention (simple maintenance) with manipulation (updating or reordering) is particularly informative, as manipulation tasks magnify age-related differences and neural capacity constraints (Bopp & Verhaeghen, 2005; Heinzel et al., 2017). Two oscillatory rhythms are central to WM: theta (4–8 Hz), associated with coordination and recruitment of cognitive resources (Reinhart & Nguyen, 2019; Roux & Uhlhaas, 2014), and alpha (8–12 Hz), linked to maintaining relevant information and inhibiting distraction (Borghini et al., 2018; Chen et al., 2023; Jensen & Mazaheri, 2010). Both are modulated by cognitive load and task phase, making them robust markers of neural efficiency and compensation across the lifespan (Klimesch, 1999; Riddle et al., 2020; Trammell et al., 2017).

In younger adults, increased frontal theta power typically supports encoding under high demands (Muthukrishnan et al., 2020; Proskovec et al., 2019a). Findings for later phases are mixed: some studies report theta increases during maintenance (Fernández et al., 2021; Ratcliffe et al., 2022), whereas others suggest that effects are largely restricted to encoding, with maintenance-related theta potentially reflecting sustained encoding activity rather than distinct processes (Lara & Wallis, 2014; Proskovec et al., 2019a; Sreenivasan et al., 2014).

Theta power is generally greater during manipulation than retention tasks and is stronger in correct trials, indicating its role in updating and reordering information (Enriquez-Geppert et al., 2014; Itthipuripat et al., 2013; Kawasaki et al., 2014).

Ageing shifts these dynamics. Under moderate demands, older adults sometimes exhibit increased frontal theta, consistent with compensatory recruitment (Festini et al., 2018; Park & Reuter-Lorenz, 2009), but when capacity is exceeded, theta responses are often attenuated compared to younger adults (Hogan et al., 2003; Kardos et al., 2014; McEvoy et al., 2001). The strongest age differences emerge in manipulation tasks, where capacity limits are taxed most (Billig et al., 2020; Bopp & Verhaeghen, 2005; Chikhi et al., 2022; D’Esposito et al., 1999).

By contrast, alpha modulation shows a more complex pattern. Reductions in alpha power, especially in frontal and posterior regions, are often linked to better performance under high demands, reflecting increased cortical excitability for updating and retrieval (Klimesch, 1997, 1999; Proskovec et al., 2016, 2019b; Scharinger et al., 2017). Conversely, increases in alpha during maintenance may facilitate inhibition of irrelevant stimuli and protection of stored information (Bonnefond & Jensen, 2012; Jensen & Mazaheri, 2010). This dual role aligns with “gating” models, in which alpha decreases facilitate encoding and updating, whereas alpha za et al., 2014). Lower alpha (∼8–increases prioritise retention (D’Ardenne et al., 2012; Man 10 Hz) has been linked to attentional updating, while upper alpha (∼10–13 Hz) supports inhibitory control and retention (Michels et al., 2008; Pavlov & Kotchoubey, 2022).

In ageing, alpha modulation does not simply weaken but becomes more variable in timing and regional distribution. Some studies report stronger alpha suppression in older adults (Springer et al., 2023), others attenuated or absent effects (McEvoy et al., 2001; Vaden et al., 2012). More nuanced patterns suggest compensatory alpha decreases during early maintenance, especially under high load, with a shift from prefrontal to posterior engagement as recall approaches (Proskovec et al., 2016; Sghirripa et al., 2021).

Taken together, theta and alpha power dynamics during maintenance and recall appear to reflect complementary strategies for balancing retention and updating across the lifespan.

However, inconsistencies in the literature highlight the need to clarify how neural efficiency, task complexity, and compensatory mechanisms interact in ageing.

The present study examined behavioural performance and oscillatory dynamics in younger and older adults during maintenance and retrieval in verbal (Letter Span) and visuospatial (Corsi Block) tasks, under both retention and manipulation conditions, using electroencephalography (EEG).

We hypothesised:

1. Behaviourally, older adults would perform worse than younger adults, particularly in manipulation and visuospatial (Corsi) tasks, where age effects are strongest (Myerson et al., 1999).
2. Theta dynamics: younger adults would show sustained increases in frontal theta across WM phases, whereas older adults would show reduced or plateaued increases, especially between encoding and maintenance/manipulation.
3. Alpha dynamics: both groups would show decreases in frontal and posterior alpha, with steeper suppression in older adults. However, if task demands exceeded capacity, older adults were expected to show attenuated or flattened modulation, reflecting protective strategies.
4. Performance–oscillation associations: task performance would correlate with theta and alpha modulation, reflecting the neural mechanisms underpinning compensation.

## METHOD

### Participants

31 healthy younger adults (18 women, 13 men, M = 21.32, SD = 3.31, range = 18 – 29 years) and 36 healthy older adults (23 women, 13 men, M = 68.69, SD = 4.94, range = 60 – 80 years) participated in the study. Younger participants were recruited through the SONA systems and public announcements at Edge Hill University, while older participants were drawn from the local community through public announcements. Exclusion criteria included left-handedness, a history of psychiatric or neurological disorders (including epilepsy and seizures), a history of alcohol or drug abuse, vision and/or hearing impairments, previous surgical procedures involving the head or spinal cord, or taking medications that might influence cognitive abilities (e.g., those causing reduced attention, fatigue, or memory impairment).

Older participants were first screened via telephone to ensure they did not meet any of the exclusion criteria. Eligible older participants were invited to take part in a study involving cognitive training and transcranial direct current stimulation (tDCS) but they were also asked for their consent to utilise their data for the present study. All older participants who provided consent and were found eligible were invited to an introductory session where their eligibility was further confirmed using the Montreal Cognitive Assessment (MoCA), the Geriatric Depression Scale (GDS) and a medical screening questionnaire that included detailed review of their clinical history. At this stage, participants were excluded if they scored below 26 points on the MoCA, more than five points on the GDS, or met any other exclusion criteria from the medical screening.

Younger participants were assessed with the Beck’s Depression Inventory (BDI) and the medical screening questionnaire, and they were excluded from the study if they scored higher than 16 points on the BDI or met any other exclusion criteria from the medical screening.

There were no significant differences between age groups in the levels of education (*t*(65) = 1.66, *p* = .10), all participants reported sleeping for at least six hours the previous night, refrained from alcohol or drug use within eight hours prior, and abstained from coffee consumption within the two hours before testing. A complete description of the study instructions and an exhaustive explanation of the functioning of EEG was provided to all participants before the beginning of the session. All participants signed a written consent. This study was approved by the Ethics Research Committee Panel of Edge Hill University and the School of Psychology.

### Working Memory tasks

#### Letter Span

Two WM tasks were employed to measure near-transfer effects, both presented in PsychoPy Software (Peirce, 2007). One of the tasks was an adapted version of the Letter Span (Figure 1A), used in previous studies (e.g., Pavlov & Kotchoubey, 2017; Postle et al., 2006; Scheeringa et al., 2009), but in the auditory modality. Each trial began with a fixation cross for one second, followed by a sequence of six consonants presented auditorily at a rate of one per second. The potential consonants included C, F, H, J, L, N, K, P, Q, R, V, W.

**Figure 1.**
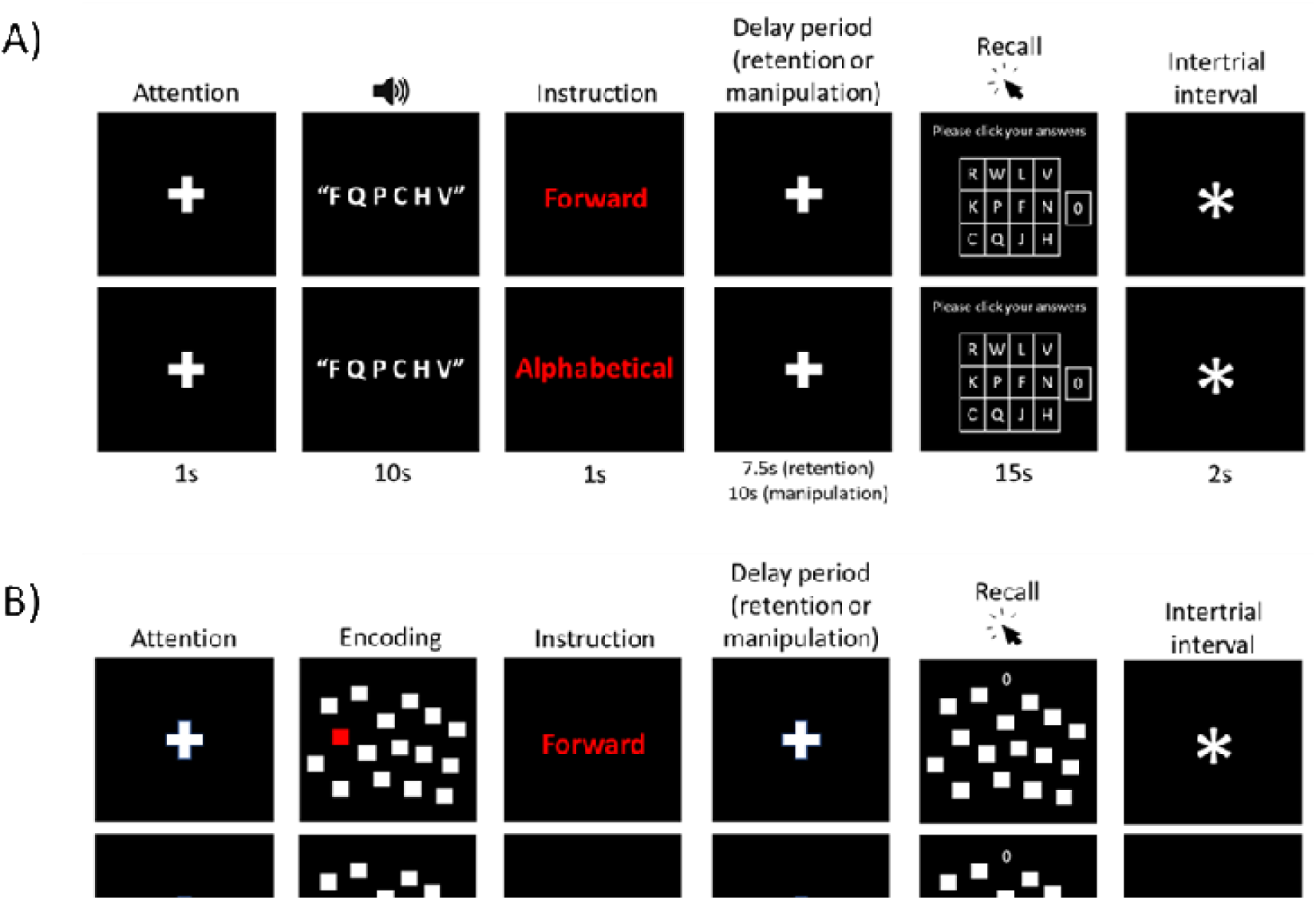
A) Adapted version of the Letter Span. Participants heard a sequence of six letters presented one by one, followed by an instruction cue indicating the recall order. After a delay period, they selected the letters from a grid in the specified order. B) Adapted version of the Corsi Block-Tapping Test. Participants saw a sequence of six squares highlighted one by one, followed by an instruction cue indicating the recall order. After a delay period, they selected the squares from the display in the specified order.

After the final letter, participants saw an instruction cue in red font, either “Forward” (retention condition) or “Alphabetical” (manipulation condition), followed by a fixation cross marking the onset of delay period. The delay lasted 7.5 seconds for the retention condition and 10 seconds for the manipulation condition. In the retention condition, participants recalled the sequence in the original order; in the manipulation condition, they reordered the sequence alphabetically before recall.

The recall phase began with a 4 × 3 grid on the screen displaying all possible letters and a zero key. Participants clicked letters with the mouse in the instructed order within 15 seconds. A two-second intertrial interval preceded the next trial.

#### Corsi Block-Tapping test

The second task was an adapted version of the Corsi Block-Tapping Test (Figure 1B) that followed a design similar to the adapted Letter Span. In this case, each trial began with sixteen white squares displayed on the screen, six of which were sequentially coloured in red for approximately one second each.

After the final square was coloured, an instruction appeared in red font showing either “Forward” (retention condition) or “Backward” (manipulation condition), followed by a fixation cross marking the onset of the delay period. The delay lasted 7.5 seconds for the retention condition and 10 seconds for the manipulation condition. In the retention condition, participants recalled the sequence of squares in the original order, and in the manipulation condition, they recalled the sequence in reversed order.

The recall phase presented the same white squares on the screen with a zero key at the top centre. Participants clicked the squares with the mouse in the instructed order within 15 seconds. They could use the zero key once per trial to fill a forgotten position. A two-second intertrial interval preceded the next trial.

The task comprised eight blocks of six trials each (24 trials per condition). Participants earned one point per correctly recalled square, with a maximum of six points per trial, keeping them focused and motivated, even if they missed some squares.

The order of the WM tasks was counterbalanced across participants, and each task took approximately 30 minutes to complete. Before starting, participants completed a practice block (three trials per condition) to familiarise themselves with the task, the mouse, and the grid layout. An additional practice block was provided if needed. The total duration of the session (including the EEG set-up) was between 2h and 2.5hrs. Participants had the opportunity to take breaks both between tests and within them (between blocks). The duration of the breaks was variable and depended on the participant.

During the EEG set-up, older participants completed the short version of the Everyday Problem-Solving Test (EPT; Willis, 1996), which comprised 28 open questions related to daily living scenarios (two questions per scenario). Participants were provided with relevant information (e.g., labels or charts) to answer the questions. This test has been primarily validated with older American populations (Moore et al., 2007; Schmitter-Edgecombe et al., 2011); therefore, a reliable comparison with younger adults was not possible. However, it was used as a warm-up exercise before the main tests, and its results were not included in the analysis. The test took approximately 15–20 minutes to complete.

#### EEG data acquisition

EEG data were acquired using a 64-channel actiCAP active electrode array and an actiCHamp amplifier system (Brain Products GmbH) at a 1000Hz sampling rate. Electrodes were placed over the scalp following the international 10 – 10 system. EEG data were online referenced to the Cz electrode. Electrodes’ impedances were kept below 10 kΩ to minimise noise and ensure a better quality of the EEG signal. BrainVision Recorder Software was used to record the signal. Data were collected continuously throughout the execution of the tasks.

#### EEG data pre-processing

EEG data were preprocessed using EEGLAB (Delorme & Makeig, 2004) in MATLAB (R2023b). Data were resampled at 512 Hz and bandpass filtered between 1 and 40 Hz. Line noise at 20 Hz was removed using the *cleanline* plugin. Bad segments were identified visually, and channels were automatically flagged for rejection if they were: 1) flat for five seconds or more, 2) <80% correlated with neighbouring channels, or 3) +/- 4 SD out of amplitude range. Independent Component Analysis (ICA) was applied to remove components reflecting >80% eye or muscle activity, with final rejection decisions guided by visual inspection. Channel and component rejection criteria were based on previous pipelines (Gil Ávila et al., 2023; Pernet et al., 2021). Rejected channels were interpolated, and data were re-referenced to the average reference.

Epochs were extracted for each WM phase (encoding, delay-retention, delay-manipulation, recall-retention, recall-manipulation) without baseline correction. Only trials with ≥ 50% accuracy were included. Participants with <25% valid trials per condition were excluded from the EEG analysis of that specific condition. This resulted in the following exclusions: Letter Span – retention: 0 Young – 0 Old; manipulation: 6 Young – 2 Old; Corsi Test – retention: 1 Young – 4 Old; manipulation: 1 Young – 11 Old; and one young and one old participant were excluded from all analyses due to recording errors. Epochs were segmented into 1-s subsegments, and residual artifacts were removed by visual inspection.

Participants with <15 valid subsegments per phase in any condition were excluded from that specific analysis (no additional exclusions). The final number of participants included in the analyses were Letter Span – retention: 30 Young – 35 Old; manipulation: 24 Young – 33 Old; Corsi Test – retention: 29 Young – 31 Old; manipulation: 29 Young – 24 Old.

#### Spectral analysis

Spectral analysis was conducted using the toolbox Fieldtrip in MATLAB (Oostenveld et al., 2011). Spectral power was extracted via a Fast Fourier Transformation (FFT) with a Hanning taper focusing on frequencies from 2Hz to 40Hz. Power was then averaged across trials (channels × frequency), log-transformed, and defined relative to each participant’s individual alpha frequency (IAF; Klimesch, 1999). The theta band ranged from five Hz below the IAF to one Hz below the lower alpha limit; alpha was split into lower and upper alpha around the IAF to capture functional distinctions within the band. Participants with values beyond ±5 SD from the median were excluded to remove extreme outliers while preserving statistical power. Given the sample size, this approach ensured that only the most extreme artefacts were excluded, without unnecessarily discarding valid data. No participants were excluded based on this criterion.

#### Statistical Analysis

Statistical analysis of behavioural results was conducted using RStudio (2023). Comparisons were assessed using a 2 (age group) × 2 (test) × 2 (condition) factorial ANOVA. Post-hoc Tukey tests followed significant main or interaction effects.

EEG data were analysed using Fieldtrip in MATLAB. Given that the primary focus of the present study was to investigate the retention, manipulation, and recall phases, Kim’s (2019) network-based model framework was employed to inform the analytical approach.

Specifically, direct comparisons were conducted between distinct WM phases to more effectively capture the dynamic transitions between them. Accordingly, oscillatory changes were calculated by subtracting the power of one WM phase from the other (e.g., delay-retention minus encoding). Importantly, as task instructions were given only after the full sequence in both tasks, encoding remained identical across retention and manipulation trials.

Cluster-based permutation independent sample t-tests (10,000 iterations) were used to compare spectral power changes across WM phases between age groups. This method was used to control for multiple comparisons (Maris & Oostenveld, 2007). Effect sizes were calculated following Meyer et al. (2021), reporting upper and lower Cohen’s d bounds. In total there were six comparisons: encoding vs delay-retention, delay-retention vs recall-retention, encoding vs recall-retention, encoding vs delay-manipulation, delay-manipulation vs recallmanipulation and encoding vs recall-manipulation. Exploration of within-group power differences between WM phases was also conducted to provide additional context to the between-group analyses.

Finally, permutation-based Spearman correlations were performed in Python (Spyder, v3.10) to link neurophysiological changes with WM performance. Electrodes were grouped into regional clusters (e.g., frontal, parietal, occipital; see Supplementary Materials). Results were corrected for multiple comparisons using false discovery rate (FDR), and bootstrapped confidence intervals were estimated to improve robustness.

## RESULTS

### Behavioural results

Significant main effects of age (*F*(1,260) = 8.74, *p* < .01, η*2 =* 0.03), test (*F*(1,260) = 50.86, *p* < .001, η*2 =* 0.16) and condition (*F*(1,260) = 101.79, *p* < .001, η*2 =* 0.21) were observed. Furthermore, significant interactions emerged between age and test (*F*(1,260) = 33.30, *p* < .001, η*2 =* 0.06) and between test and condition (*F*(1,260) = 25.33, *p* < .001, η*2 =* 0.04). The three-way interaction (age × test × condition) was not significant (*F*(1,260) = 0.10, *p* = .76, η*2* < 0.01).

Post hoc analysis revealed that no significant age-group differences were found in Letter Span performance (B = 0.29, SE = 0.15, *t*(260) = 1.99, *p* = .19), whereas in the Corsi Test older adults performed significantly worse than younger adults (B = -0.90, SE = 0.15, *t*(260) = -6.17, *p* < .001, *d* = -1.10). Notably, older adults performed significantly better on the Letter Span than on the Corsi Test (B = 1.28, SE = 0.10, *t*(260) = 9.15, *p* < .001, *d* = 1.36), but younger adults showed no performance differences between the two tests (B = 0.09, SE = 0.15, *t*(260) = 0.62, *p* = .93).

Regarding the test × condition interaction, during the retention condition, participants performed significantly better in the Letter Span than in the Corsi Test (B = 1.20, SE = 0.15, *t*(260) = 8.26, *p* < .001, *d* = 1.33), but no difference between tests was observed in the manipulation condition (B = 0.17, SE = 0.15, *t*(260) = 1.19, *p* = .63). Finally, both age groups performed significantly better in the retention condition than in manipulation (Letter Span: B = 1.55, SE = 0.15, *t*(260) = 10.66, *p* < .001, *d* = 1.73; Corsi Test: B = 0.52, SE = 0.15, *t*(260) = 3.59, *p* < .01, *d* = 0.58).

### Age differences in power changes

#### Letter Span – Theta Band

From encoding to delay, younger adults showed significantly stronger theta power increases than older adults in both conditions (retention: *p* < .001, *d* = [1.14, 1.44]; manipulation: *p* < .01, *d* = [1.15, 1.30]), with effects localised in mid-frontal, central, and temporal regions (see Figure 3A). Exploration of oscillatory trends revealed that in the retention condition, younger adults exhibited significant theta increases, whereas older adults showed significant decreases. In contrast, in the manipulation condition, younger adults did not display any significant theta increases, but older adults showed again significant decreases.

**Figure 2.**
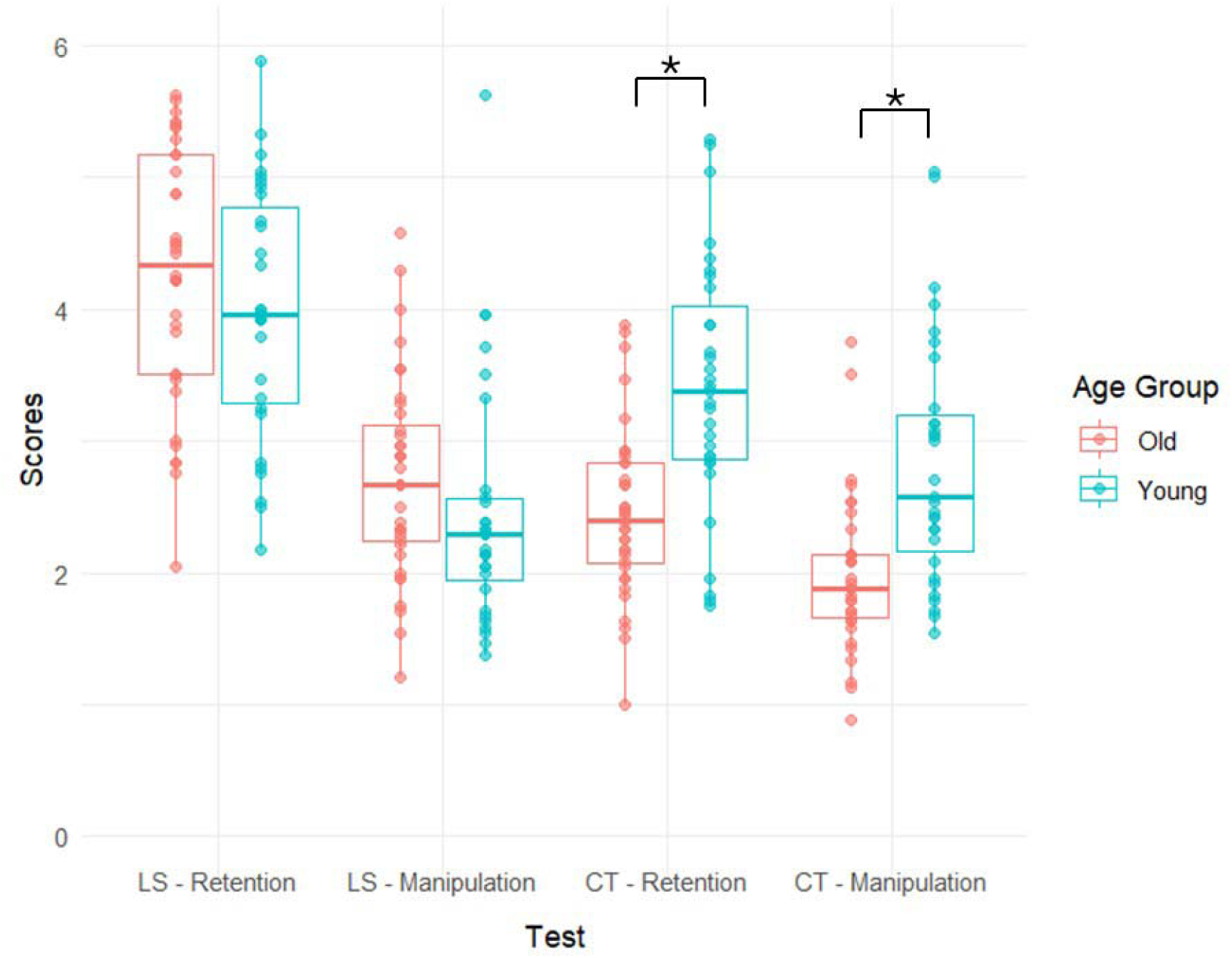
Comparison of scores of all WM tests between age groups. LS: Letter Span. CT: Corsi Test. Significant differences (*p* < 0.05) are marked with an asterisk (*). Younger adults performed significantly better than older adults in the Corsi Test. No significant differences were found in the Letter Span.

**Figure 3.**
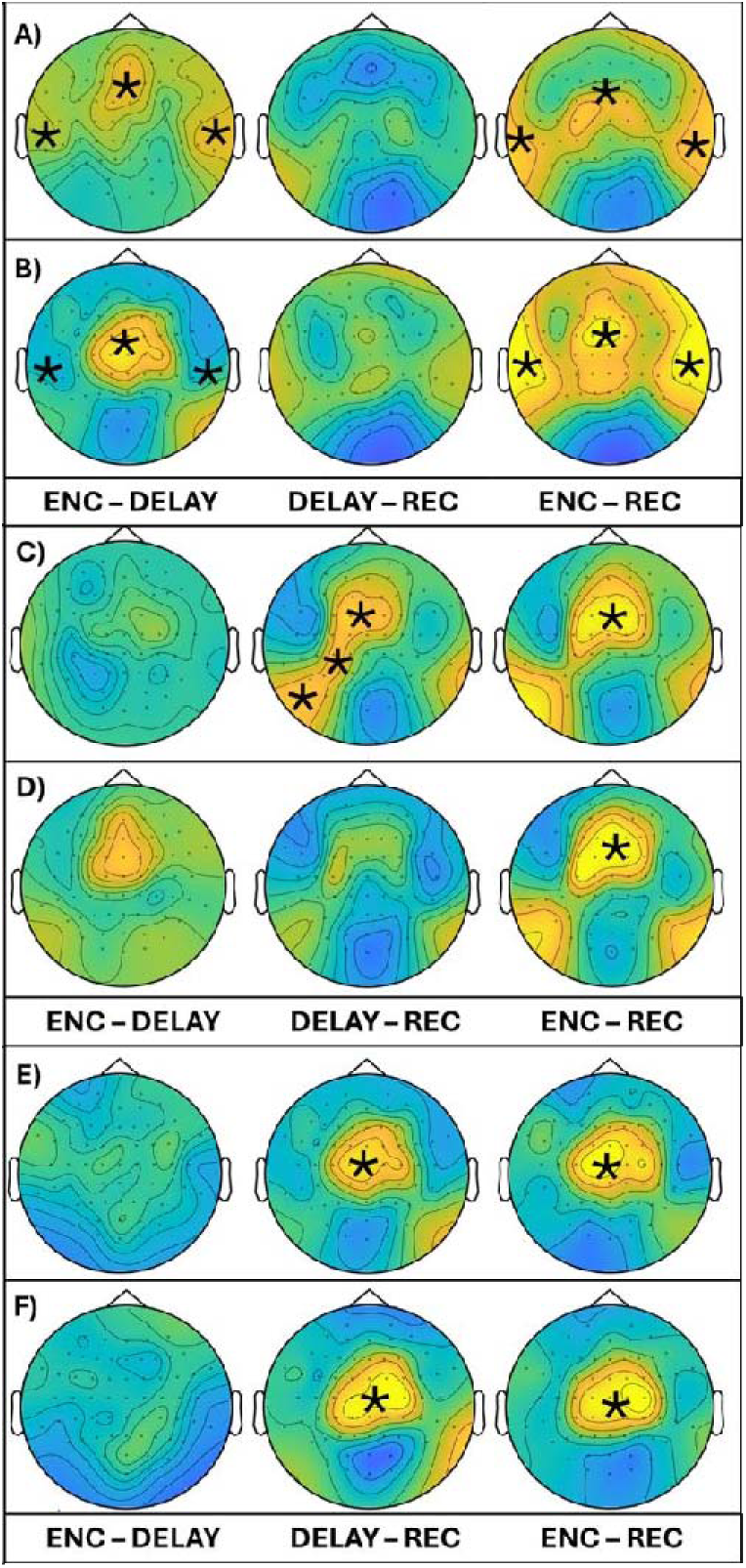
Topographies of age-group differences in power changes in the Letter Span. Significant and marginally significant differences (*p* ≤ 0.05) are marked with an asterisk (*). Colours indicate group differences: yellow = higher power in younger adults, blue = higher power in older adults. ENC = encoding; DELAY = delay period; REC = recall. A) Retention – Theta: younger adults showed significantly stronger theta power increases than older adults in mid-frontal, central, and temporal regions from encoding to delay and across the full trial (encoding to recall). B) Manipulation – Theta: younger adults showed significantly stronger theta power increases than older adults in mid-frontal, central, and temporal regions from encoding to delay and across the full trial. C) Retention – Lower alpha: older adults showed stronger lower alpha power decreases than younger adults in mid-frontal and left parietal and parieto-occipital regions from delay to recall; across the full trial, this effect was observed in the mid-frontal region. D) Manipulation – Lower alpha: older adults showed stronger lower alpha power decreases than younger adults in the mid-frontal region across the full trial. E) Retention – Upper alpha: older adults showed stronger upper alpha power decreases than younger adults in the mid-frontal region, both from delay to recall and across the full trial. F) Manipulation – Upper alpha: older adults showed stronger upper alpha power decreases than younger adults in the mid-frontal region, both from delay to recall and across the full trial.

**Figure 4.**
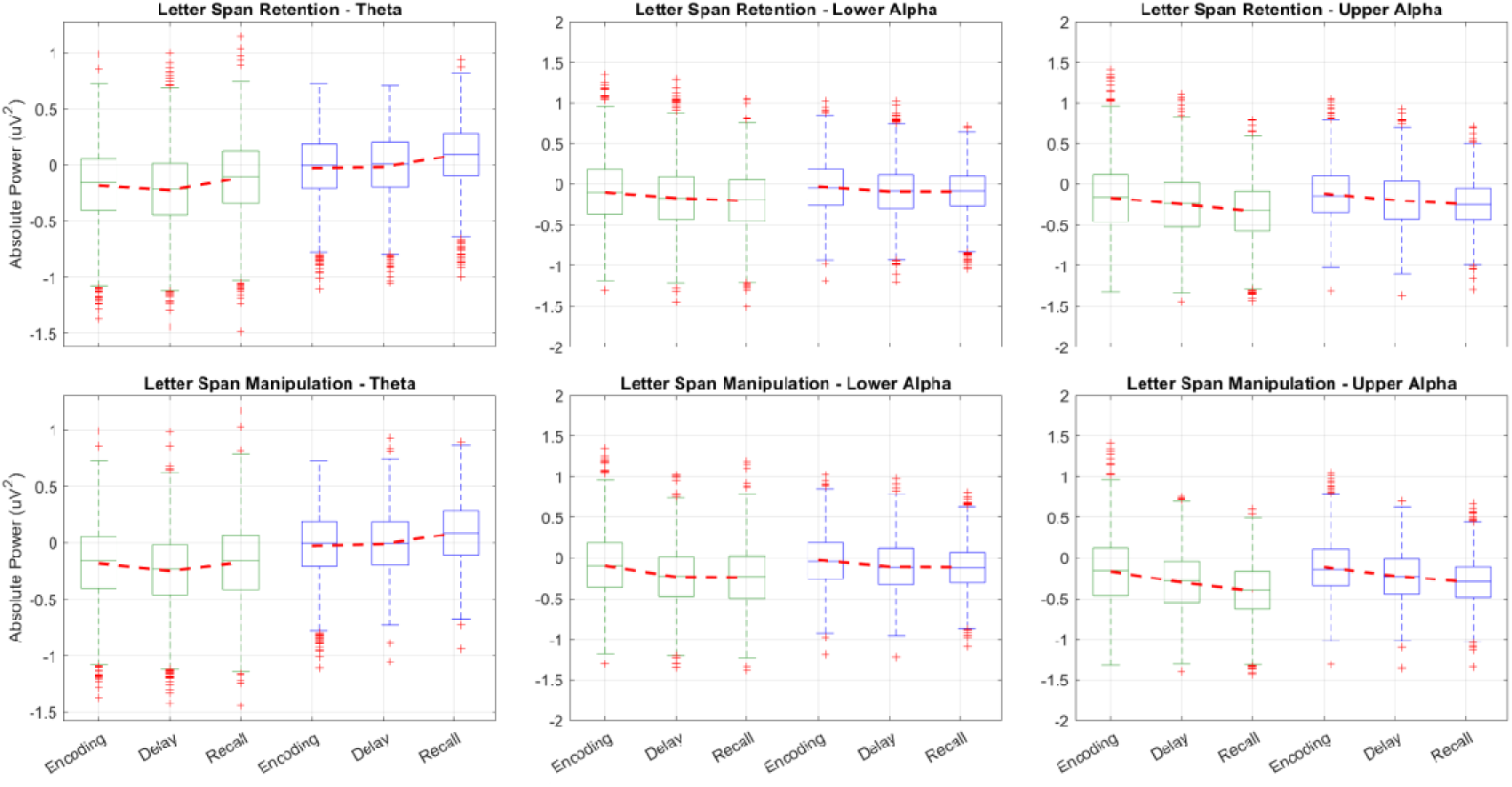
Boxplots representing differences in power changes within each age group across WM phases in the Letter Span. Green boxplots represent older participants, and blue boxplots represent younger participants.

From delay to recall, no significant age-group differences were observed in either condition. Oscillatory trends indicated that both groups exhibited significant theta increases in both conditions.

Overall, across the full trial (encoding to recall), both age groups showed significant theta power increases in both conditions, but these increases were significantly stronger in younger adults in mid-frontal and temporal regions (retention: *p* < .05, *d* = [0.79, 0.93]; manipulation: *p* < .01, *d* = [1.00, 1.15]) (see Figures 3A and 3B).

#### Letter Span – Lower Alpha Band

From encoding to delay, no significant age-group differences in lower alpha power changes were observed in either of the conditions. Exploration of oscillatory trends revealed that both age groups exhibited a significant decrease in lower alpha power in both conditions.

From delay to recall, older adults showed marginally stronger lower alpha power decreases than younger adults only in the retention condition (*p* = .05, *d* = [0.65, 0.74]), with effects localised in mid-frontal and left parietal and parieto-occipital regions (see Figure 3C).

Oscillatory trends indicated that younger adults showed no lower alpha power changes in either of the conditions, but older adults exhibited a significant decrease in both conditions.

Overall, across the full trial, both age groups showed significant decreases in lower alpha power in both conditions, but these decreases were significantly stronger in older adults in the mid frontal region (retention: *p* < .05, *d* = [0.74, 0.85]; manipulation: *p* < .05, *d* = [0.80, 0.97]) (see Figures 3C and 3D).

#### Letter Span – Upper Alpha Band

From encoding to delay, no significant age-group differences in upper alpha power changes were observed in either of the conditions. Exploration of oscillatory trends revealed that both age groups displayed a significant decrease in upper alpha power in both conditions.

From delay to recall, older adults showed significantly stronger upper alpha power decreases than younger adults in both condition (retention: *p* < .05, *d* = [1.00, 1.10]; manipulation: *p* = .05, *d* = [0.85, 0.95]), with effects localised in the mid-frontal region (see Figure 3E and 3F). Oscillatory trends indicated that both age groups showed a significant decrease in upper alpha power in both conditions.

Overall, across the full trial, both age groups showed significant decreases in upper alpha power in both conditions, but these decreases were significantly stronger in older adults in the mid-frontal region (retention: *p* = .05, *d* = [0.74, 0.81]; manipulation: *p* = .05, *d* = [0.79, 0.92]) (see Figures 3E and 3F).

Full exploratory statistics are reported in Table 1. An extended version of Table 1, including the specific brain regions where these changes occurred, is provided in the supplementary materials (Table S1).

**Table 1.**
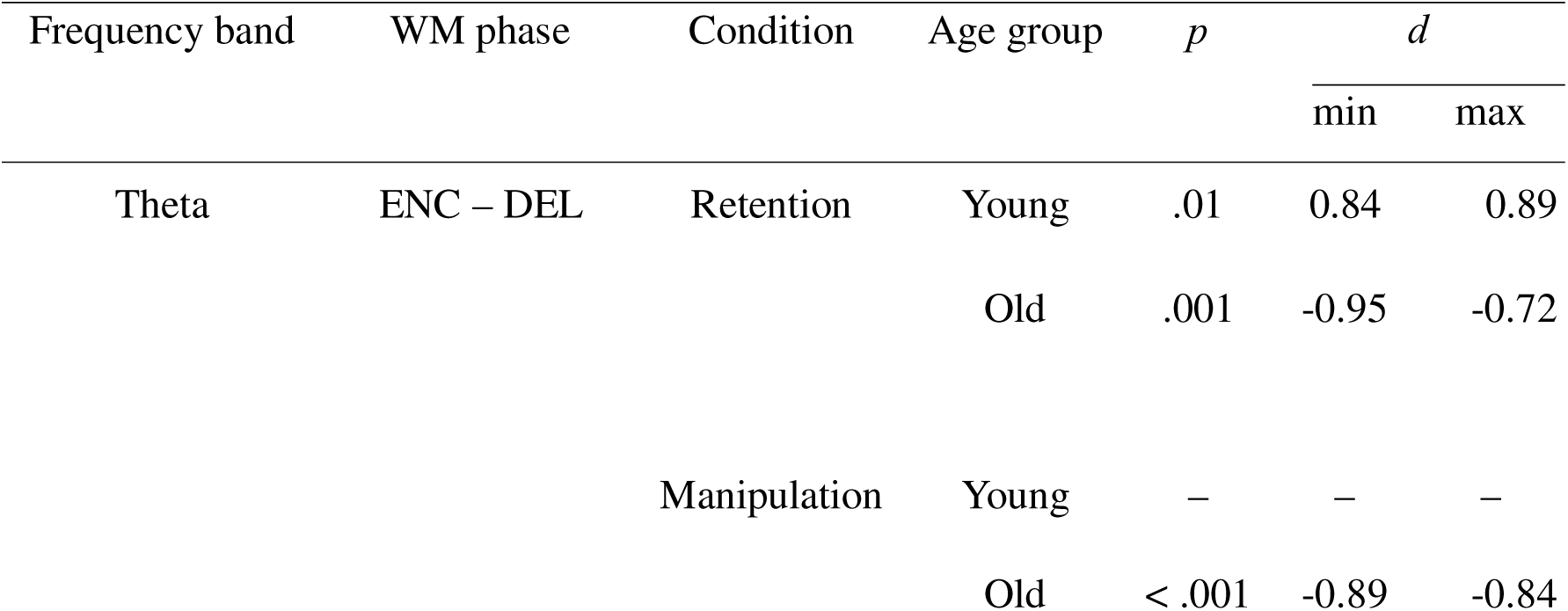
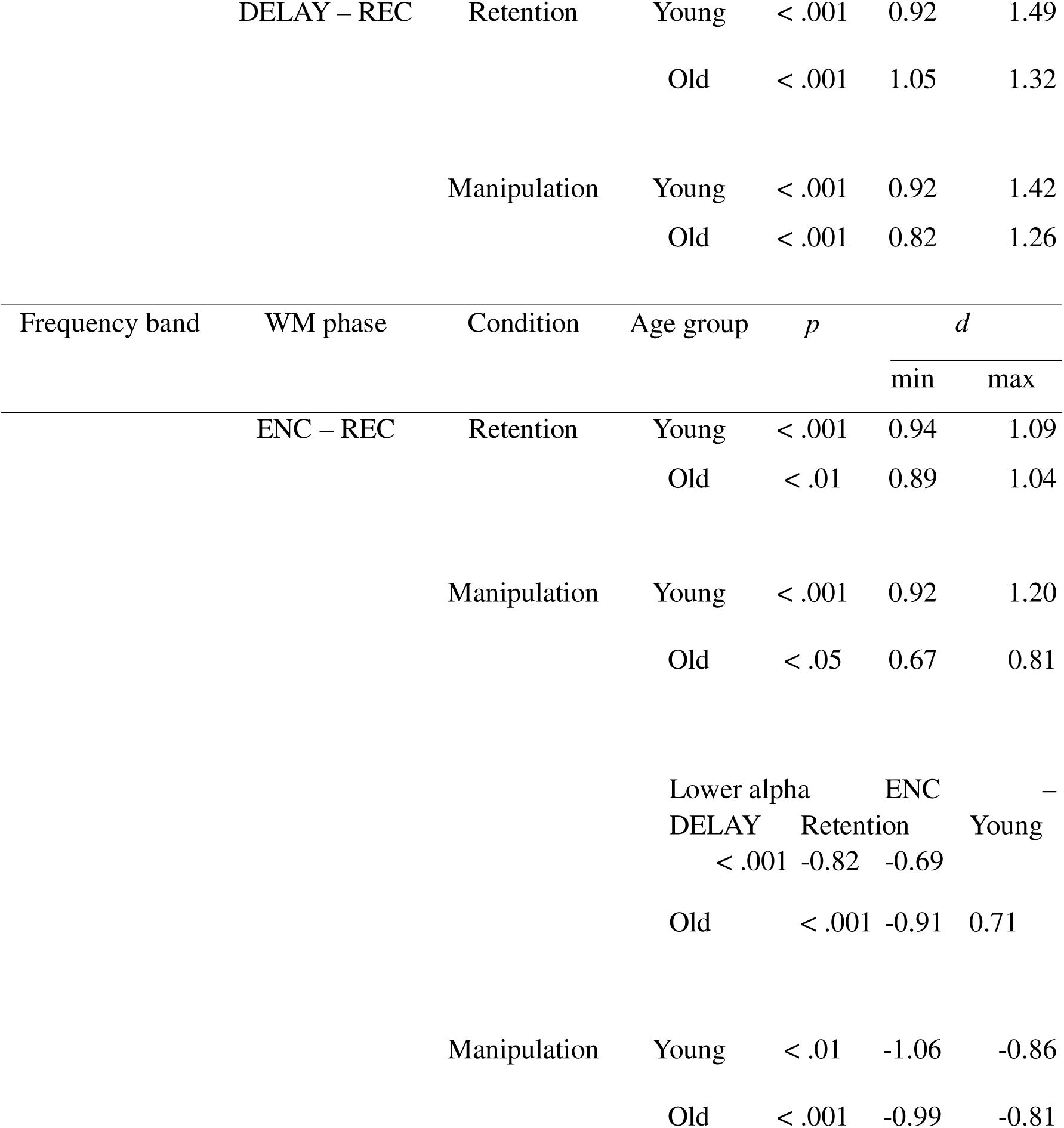
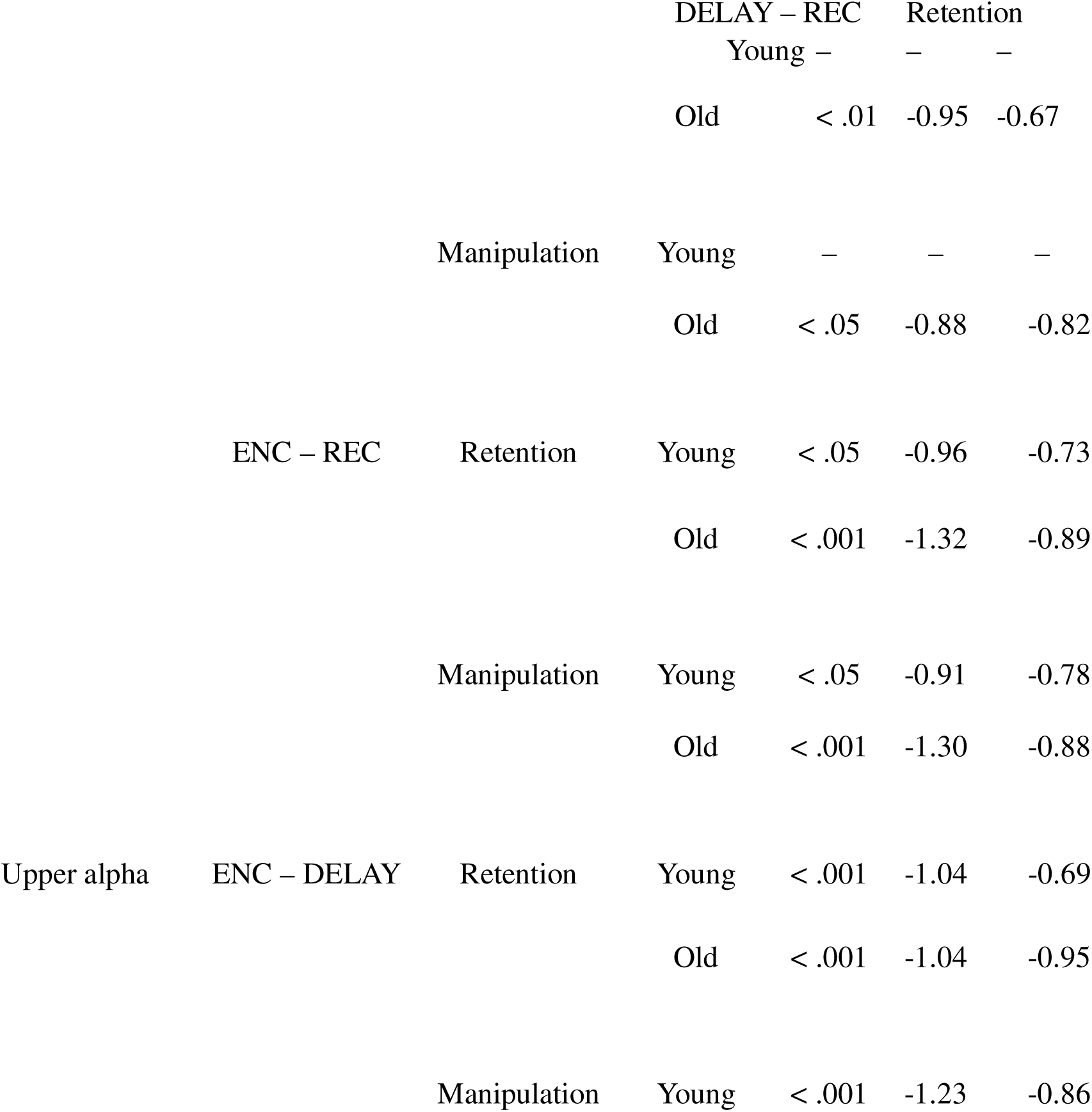
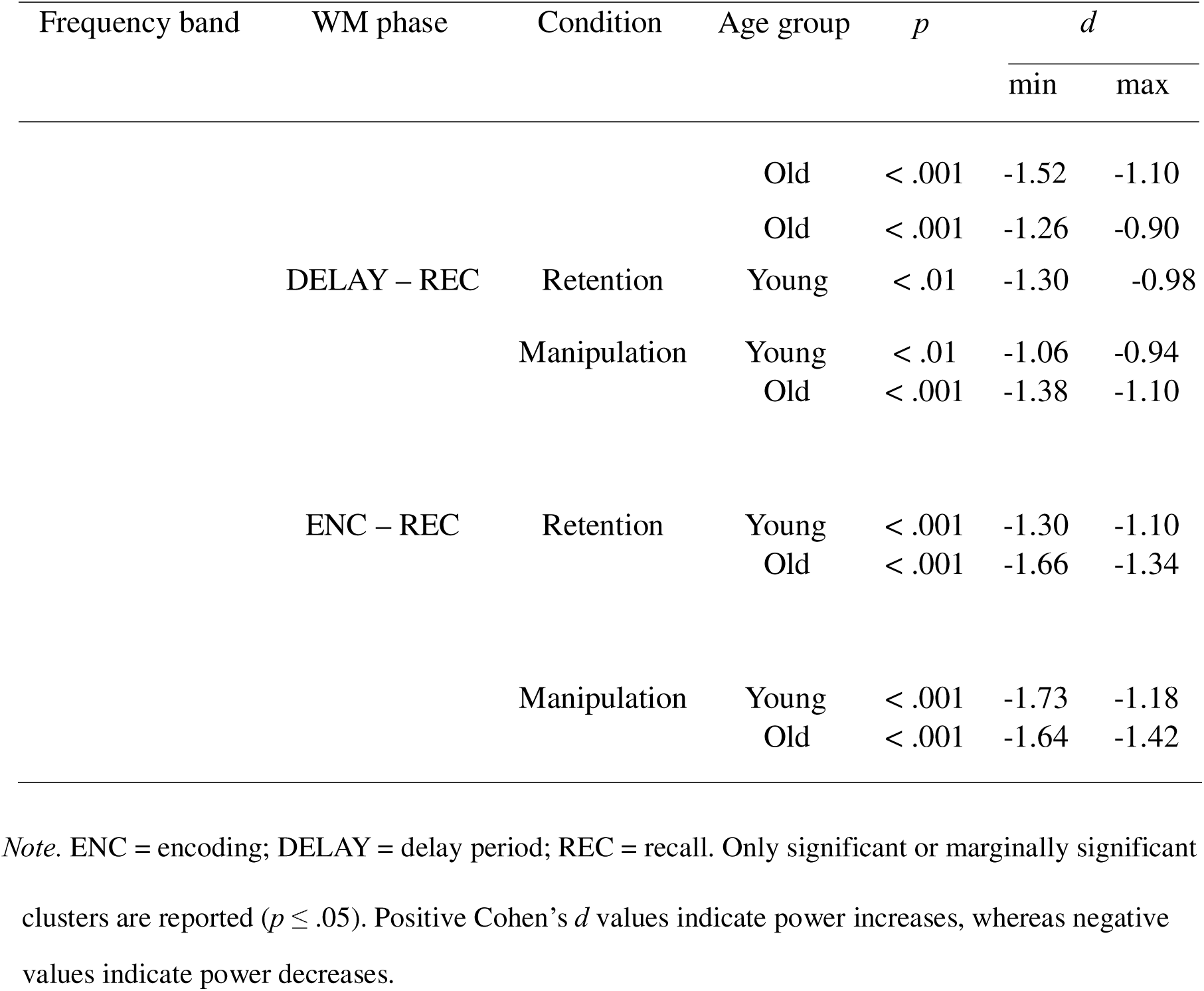
Exploration of oscillatory trends: significant differences in power changes within each age group across WM phases in the Letter Span.

#### Corsi Test – Theta Band

From encoding to delay, no significant age-group differences in theta power changes were observed in either of the conditions. Exploration of oscillatory trends revealed that both age groups exhibited significant theta power decreases in both conditions.

From delay to recall, younger adults showed significantly stronger theta power increases compared to older adults in both conditions (retention: *p* < .05, *d* = [0.85, 0.87]; manipulation: *p* = .05, *d* = [0.95, 1.05]). In the retention condition, effects were localised in the mid-frontal, central, and left temporal regions, whereas in the manipulation condition, effects were localised in the mid-frontal region (see Figures 5A and 5B). Oscillatory trends indicated that both age groups displayed significant theta power increases in both conditions.

**Figure 5.**
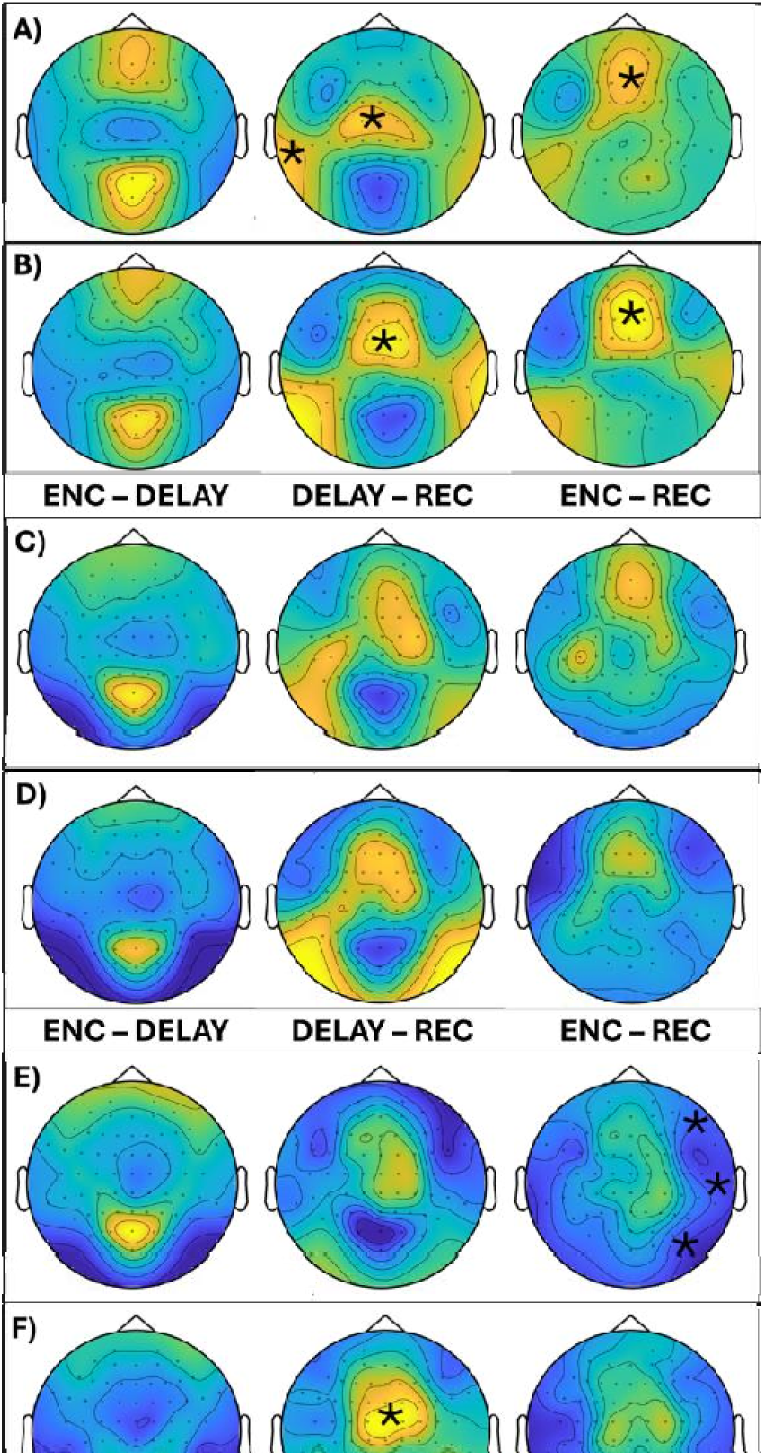
Topographies of age-group differences in power changes in the Corsi Test. Significant and marginally significant differences (p ≤ 0.05) are marked with an asterisk (*). Colours indicate group differences: yellow = higher power in younger adults, blue = higher power in older adults. ENC = encoding; DELAY = delay period; REC = recall. A) Retention – Theta: younger adults showed significantly stronger theta power increases than older adults in mid-frontal, central, and left temporal regions from encoding to delay; across the full trial (encoding to recall), this effect was observed in the mid-frontal region. B) Manipulation – Theta: younger adults showed significantly stronger theta power increases than older adults in the mid-frontal region from delay to recall and across the full trial. C) Retention – Lower alpha: no significant age-group differences in power were observed across WM phases. D) Manipulation – Lower alpha: no significant age-group differences in power were observed across WM phases. E) Retention – Upper alpha: younger adults showed stronger upper alpha power decreases than older adults in right frontal, temporal and parieto-occipital regions across the full trial. F) Manipulation – Upper alpha: older adults showed stronger upper alpha power decreases than younger adults in the mid-frontal region from delay to recall.

**Figure 6.**
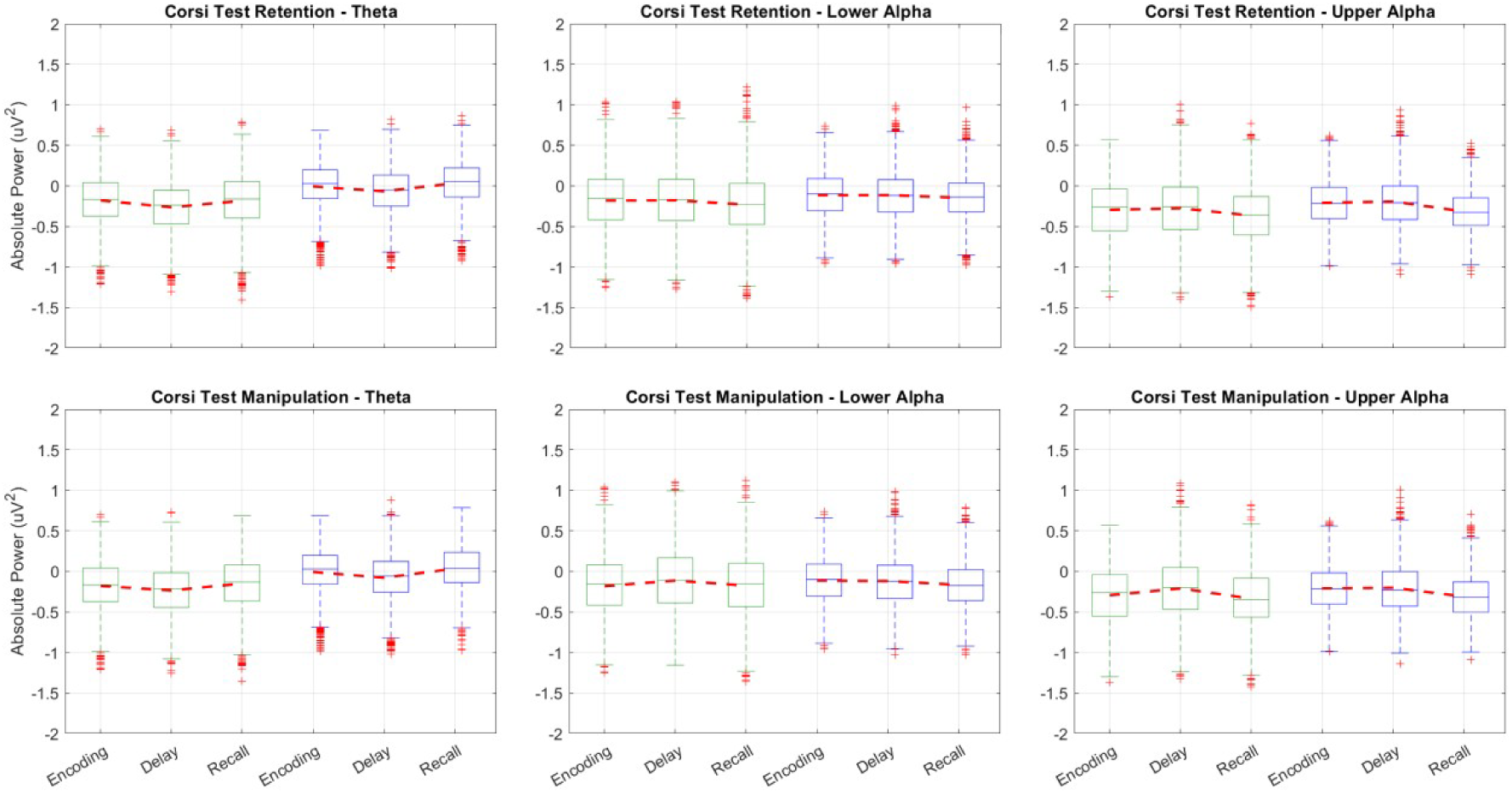
Boxplots representing differences in power changes within each age group across WM phases in the Corsi Test. Green boxplots represent older participants, and blue boxplots represent younger participants.

Overall, across the full trial, younger adults showed significantly stronger theta power increases than older adults in both conditions (retention: *p* < .05, *d* = [0.81, 0.96]; manipulation: *p* < .05, *d* = [1.00, 1.19]), with effects localised in the mid-frontal region (see Figures 5A and 5B). Oscillatory trends indicated that younger, but not older adults, increased theta power significantly across phases.

#### Corsi Test – Lower Alpha Band

No significant age-group differences in lower alpha power changes were observed across WM phases. Exploration of oscillatory trends revealed that, from delay to recall and across the full trial, both age groups exhibited significant lower alpha power decreases in both conditions.

#### Corsi Test – Upper Alpha Band

From encoding to delay, no significant age-group differences in upper alpha power changes were observed in either of the conditions. Exploration of oscillatory trends revealed that both age groups showed a significant increase in upper alpha power in both conditions.

From delay to recall, older adults displayed a significantly stronger upper alpha power decrease compared to younger adults only in the manipulation condition (*p* < .05, *d* = [0.96, 0.99]), with effects localised in the mid-frontal region (see Figures 5F). Oscillatory trends indicated that both age groups exhibited a significant decrease in power in both conditions.

Overall, across the full trial, both age groups showed significant decreases in upper alpha power in both conditions, but the decrease in the retention condition was significantly stronger in younger adults in right frontal, temporal and parieto-occipital regions (*p* < .05, *d* = [-0.94, -0.89]) (see Figures 5E).

Full exploratory statistics are reported in Table 2. An extended version of Table 2, including the specific brain regions where these changes occurred, is provided in the supplementary materials (Table S2).

**Table 2.**
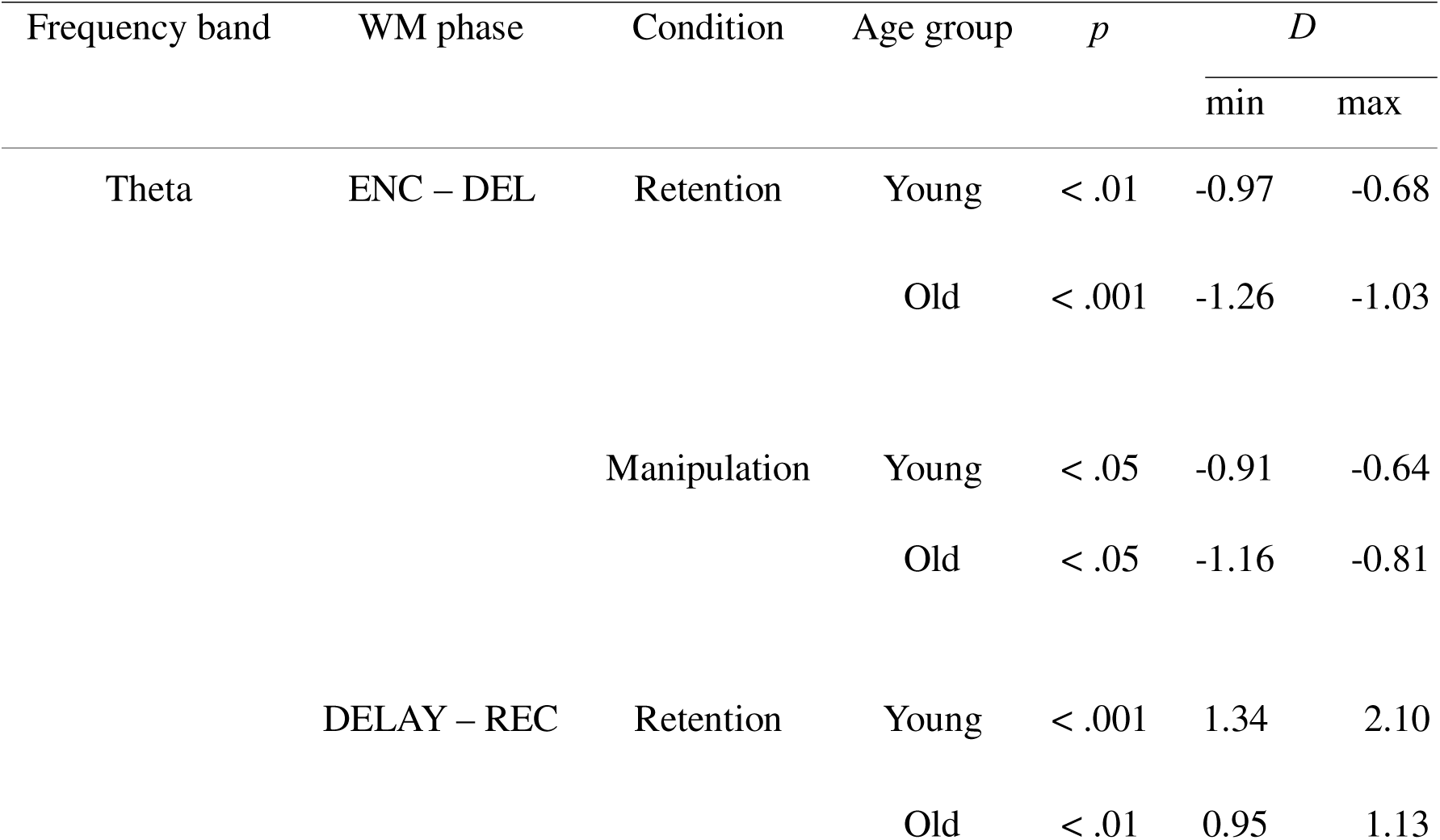
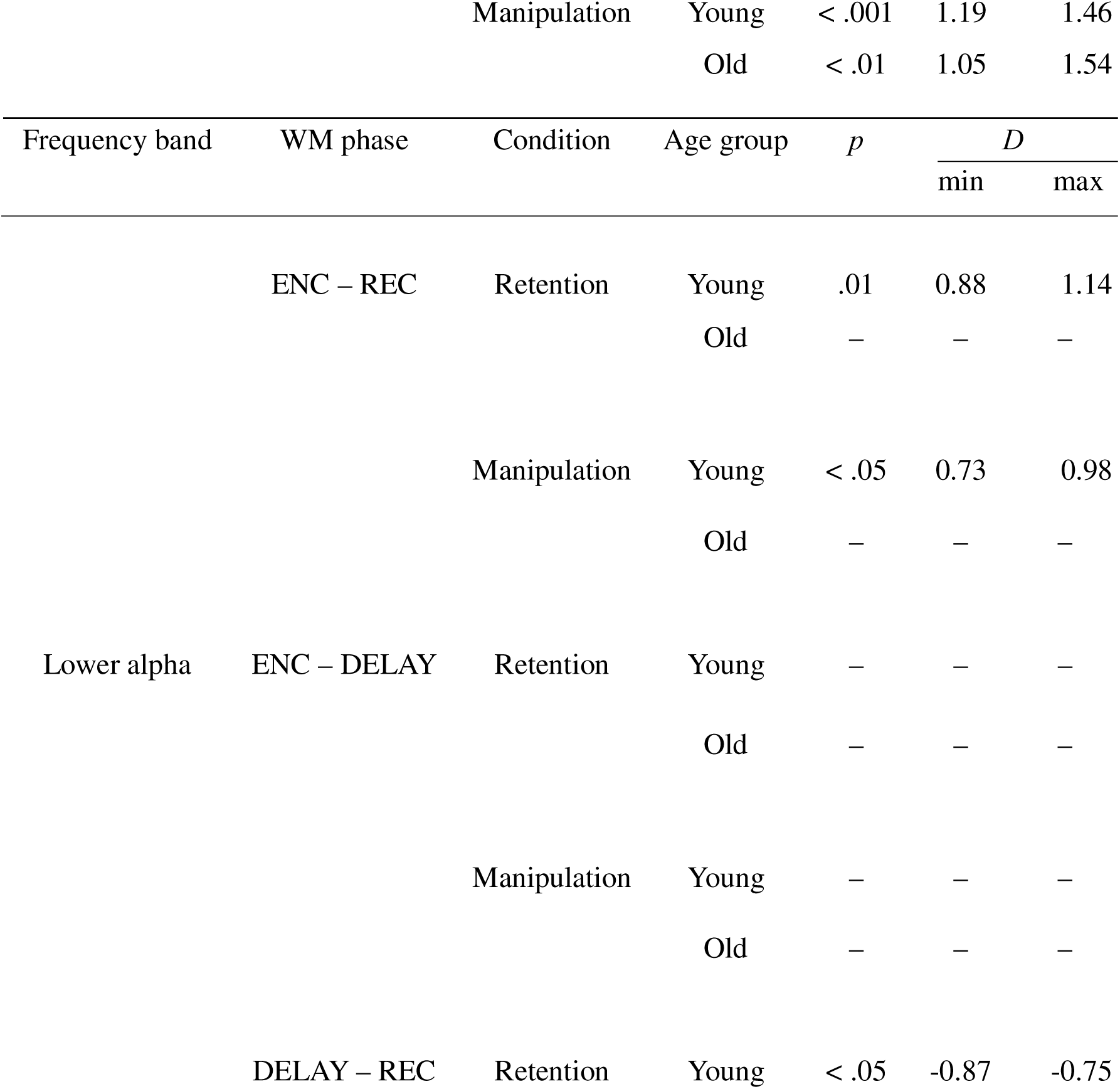
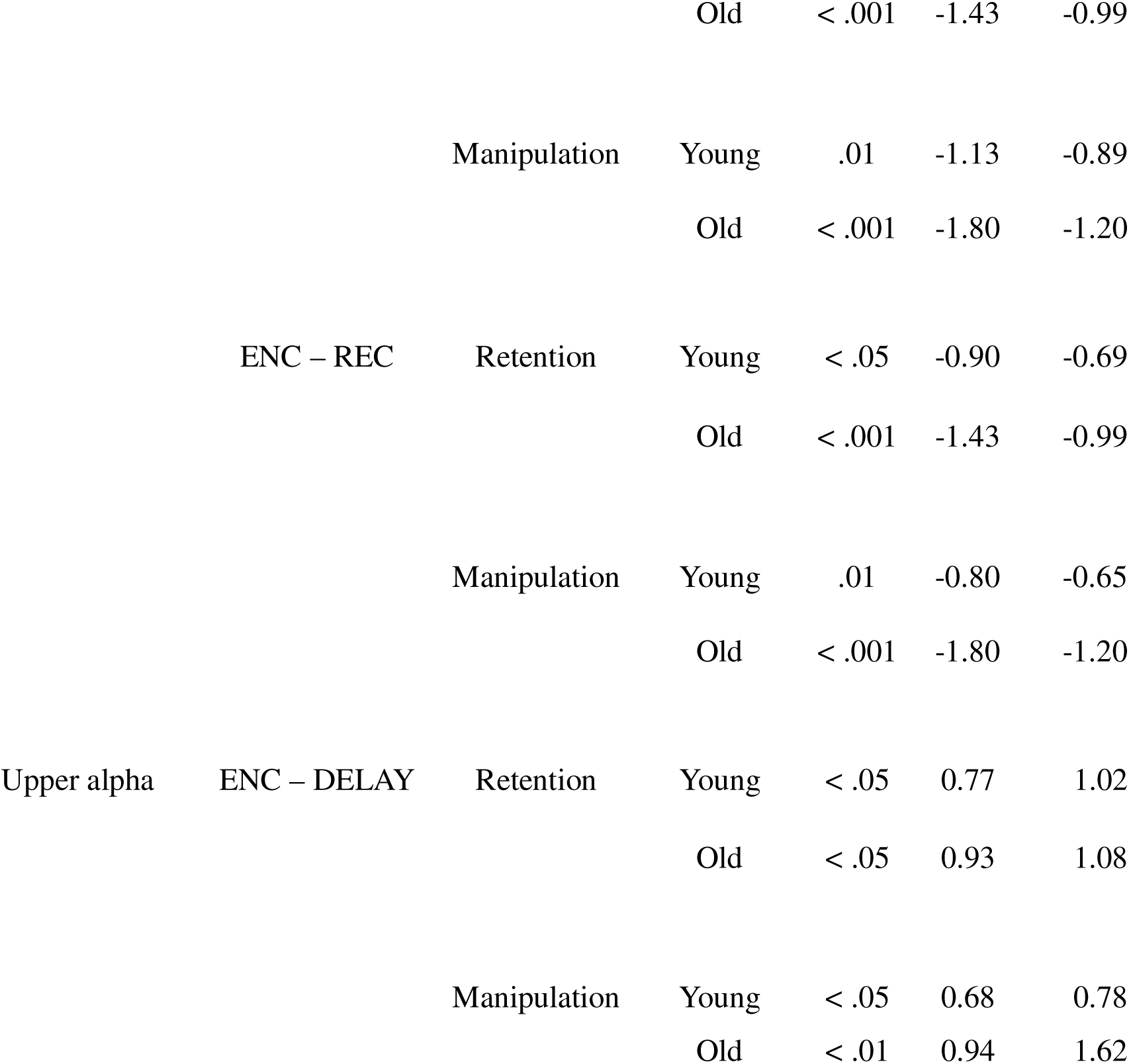
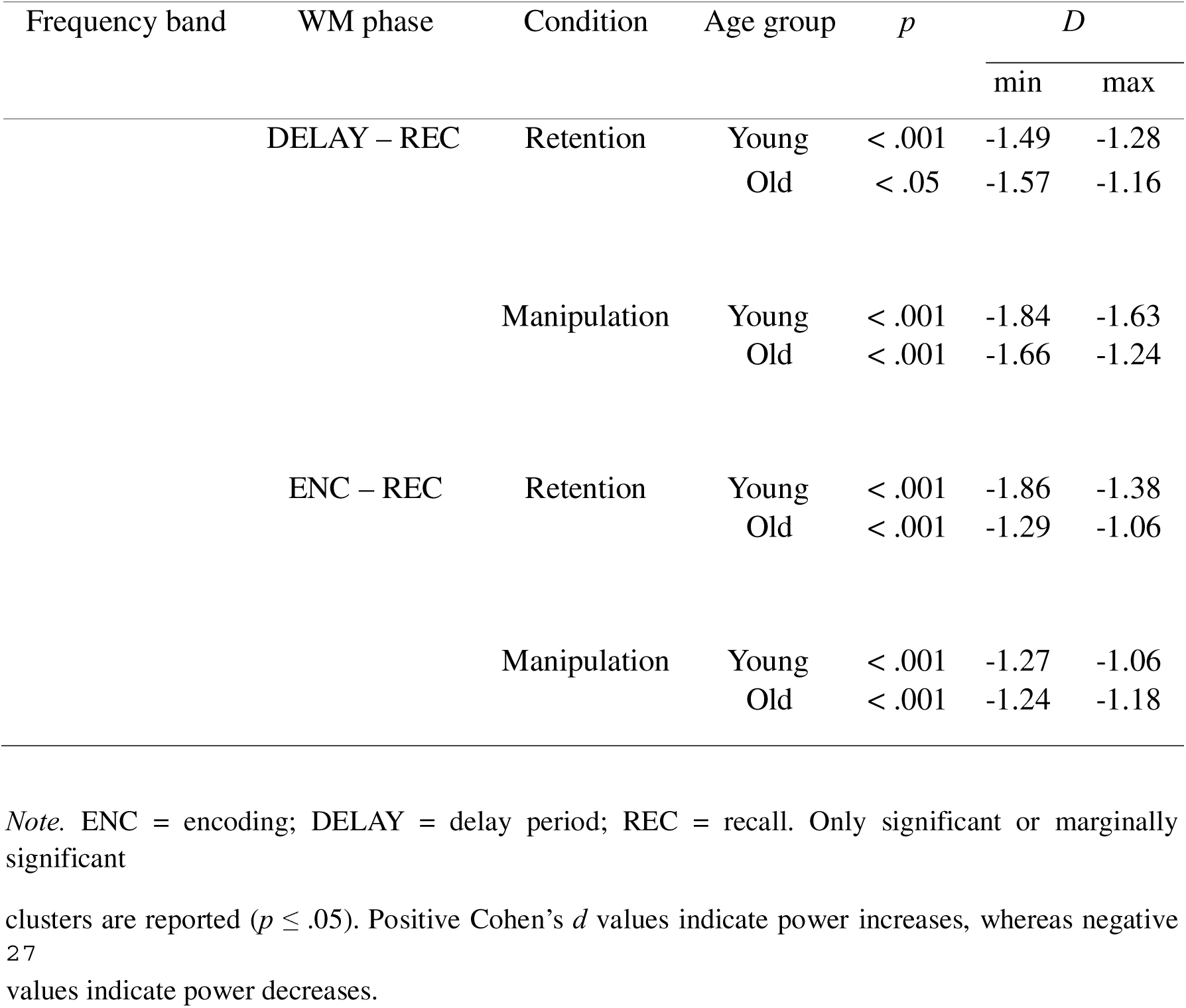
Exploration of oscillatory trends: significant differences in power changes within each age group across WM phases in the Corsi Test.

| Frequency band | WM phase | Condition | Age group | $p$ | $D$ | |
| --- | --- | --- | --- | --- | --- | --- |
|  |  |  |  |  | min | max |
| Theta | ENC – DEL | Retention | Young | < .01 | -0.97 | -0.68 |
|  |  |  | Old | < .001 | -1.26 | -1.03 |
|  |  | Manipulation | Young | < .05 | -0.91 | -0.64 |
|  |  |  | Old | < .05 | -1.16 | -0.81 |
|  | DELAY – REC | Retention | Young | < .001 | 1.34 | 2.10 |
|  |  |  | Old | < .01 | 0.95 | 1.13 |

|  |  | Manipulation | Young | < .001 | 1.19 | 1.46 |
| --- | --- | --- | --- | --- | --- | --- |
|  |  |  | Old | < .01 | 1.05 | 1.54 |
| Frequency band | WM phase | Condition | Age group | *p* | *D* |  |
|  |  |  |  |  | min | max |
|  | ENC – REC | Retention | Young | .01 | 0.88 | 1.14 |
|  |  |  | Old | – | – | – |
|  |  | Manipulation | Young | < .05 | 0.73 | 0.98 |
|  |  |  | Old | – | – | – |
| Lower alpha | ENC – DELAY | Retention | Young | – | – | – |
|  |  |  | Old | – | – | – |
|  |  | Manipulation | Young | – | – | – |
|  |  |  | Old | – | – | – |
|  | DELAY – REC | Retention | Young | < .05 | -0.87 | -0.75 |
|  |  |  | Old | < .001 | -1.43 | -0.99 |
|  |  | Manipulation | Young | .01 | -1.13 | -0.89 |
|  |  |  | Old | < .001 | -1.80 | -1.20 |
|  | ENC – REC | Retention | Young | < .05 | -0.90 | -0.69 |
|  |  |  | Old | < .001 | -1.43 | -0.99 |
|  |  | Manipulation | Young | .01 | -0.80 | -0.65 |
|  |  |  | Old | < .001 | -1.80 | -1.20 |
| Upper alpha | ENC – DELAY | Retention | Young | < .05 | 0.77 | 1.02 |
|  |  |  | Old | < .05 | 0.93 | 1.08 |
|  |  | Manipulation | Young | < .05 | 0.68 | 0.78 |
|  |  |  | Old | < .01 | 0.94 | 1.62 |
|  |  |  |  |  | min | max |
|  | DELAY – REC | Retention | Young | < .001 | -1.49 | -1.28 |
|  |  |  | Old | < .05 | -1.57 | -1.16 |
|  |  | Manipulation | Young | < .001 | -1.84 | -1.63 |
|  |  |  | Old | < .001 | -1.66 | -1.24 |
|  | ENC – REC | Retention | Young | < .001 | -1.86 | -1.38 |
|  |  |  | Old | < .001 | -1.29 | -1.06 |
|  |  | Manipulation | Young | < .001 | -1.27 | -1.06 |
|  |  |  | Old | < .001 | -1.24 | -1.18 |

*Note.* ENC = encoding; DELAY = delay period; REC = recall. Only significant or marginally significant
clusters are reported ( $p \leq .05$ ). Positive Cohen's $d$ values indicate power increases, whereas negative values indicate power decreases.

### Correlations between task performance and power changes

#### Letter Span – Retention condition

No significant correlations between task performance and power changes were found in either age group.

#### Letter Span – Manipulation condition

In younger adults, task performance was positively correlated with theta power changes from encoding to recall in several regions: parietal (left: *r* = .60, 95% CI [.21, .86], *p* < .001, FDR-corrected; right: *r* = .51, 95% CI [.15, .73], *p* < .05, FDR-corrected), right frontal (*r* = .60, 95% CI [.24, .82], *p* = .01, FDR-corrected), and left temporal (*r* = .48, 95% CI [.01, .79], *p* < .05, FDR-corrected). These findings indicate that stronger theta power increases in younger adults were associated with better performance.

Regarding older adults, task performance was negatively correlated with lower alpha power changes from encoding to delay in right parietal regions (*r* = -.54, 95% CI [-.76, -.24], *p* < .001, FDR-corrected). These findings indicate that stronger lower alpha power decreases in older adults were associated with better performance.

#### Corsi Test – Retention condition

In younger adults, task performance was positively correlated with theta power changes from delay to recall in left temporal regions (*r* = .57, 95% CI [.25, .80], *p* < .05, FDR-corrected). These findings indicate that stronger theta power increases in younger adults were associated with better performance.

Regarding older adults, task performance was negatively correlated with upper alpha power changes from encoding to delay in right parietal regions (*r* = .56, 95% CI [.23, .78], *p* < .05, FDR-corrected). These findings indicate that stronger upper alpha power increases in older adults were associated with better performance.

#### Corsi Test – Manipulation condition

No significant correlations between task performance and power changes were found in either age group.

## DISCUSSION

In the present study, we examined age-related differences in oscillatory dynamics during high cognitive load working memory (WM) tasks, integrating behavioural performance with electroencephalography (EEG) markers across verbal and visuospatial domains. Our investigation was guided by four a priori hypotheses: (1) older adults would perform significantly worse than younger adults, with the largest deficits expected in manipulation conditions and in the visuospatial Corsi Test; (2) younger adults would exhibit sustained increases in frontal theta power across WM phases, whereas older adults would show plateaued or reduced increases, particularly between encoding and the delay interval; (3) both groups would demonstrate continuous decreases in frontal and posterior alpha power, with a steeper decline in older adults—or, if capacity limits were exceeded, a flattening of alpha modulation in the older group; and (4) changes in theta and alpha would correlate with trial-by-trial performance indices, thereby revealing putative compensatory mechanisms.

### Behavioural Results

As anticipated, younger adults outperformed older adults on the Corsi Test. This pattern aligns with extensive evidence that visuospatial WM and spatial abilities show marked age-related decline, reflecting vulnerabilities in both short-term storage and manipulation of spatial representations (Borella et al., 2014; Briggs et al., 1999; Brown, 2016; Carlesimo et al., 1994; Myerson et al., 1999; Salthouse, 2010). In the context of the present paradigm, the Corsi demands likely compounded processing and maintenance costs under substantial load, thereby exposing capacity constraints more readily in older participants. By contrast, the absence of age-related differences in the Letter Span suggests that the verbal task demands remained within the compensatory bandwidth of older adults, enabling the deployment of effective strategies (e.g., rehearsal, chunking, task-familiar routines) sufficient to sustain performance at levels comparable to younger adults (Chikhi et al., 2022; Graves, 2018; Klencklen et al., 2017).

Unexpectedly, there were no significant age-group differences across the retention and manipulation conditions when collapsed across modality. Both groups exhibited poorer performance in manipulation relative to retention, consistent with literature showing that manipulation engages more complex, multicomponent processes (e.g., updating, reordering, interference resolution) and requires flexible shifts in strategy as well as broader neural recruitment (D’Esposito et al., 1999, 2000; Itthipuripat et al., 2013; Román-López et al., 2019).

The uniform decline in both groups under manipulation may have reduced between-group variance, masking age effects at the condition level despite clear modality-specific differences. Thus, while the change in sensory modality disproportionately challenged older adults (Letter Span vs Corsi), the change in WM operation (retention vs manipulation) exerted a robust, group-general cost. Together, these outcomes support the view that older adults can maintain verbal WM performance through compensatory strategies when demands are moderate, but reach neural capacity limits earlier in visuospatial contexts. Notably, when task complexity imposes broad constraints—as in manipulation—age disparities may be attenuated because both younger and older adults operate closer to their respective capacity ceilings.

### Neurophysiological Results

#### Theta Power Changes

In line with our hypotheses, younger adults showed stronger theta power increases than older adults, with effects varying by WM phase and region. During the Letter Span, younger adults exhibited significantly greater theta augmentation from encoding to delay and across the entire trial (encoding to recall) under both retention and manipulation, with effects localised chiefly to mid-frontal and temporal cortices. In the Corsi Test, a similar age advantage emerged, but the most pronounced differences occurred from delay to recall and across the full epoch, with a mid-frontal focus and additional left temporal involvement in the retention condition.

These patterns indicate that younger adults more effectively mobilised and sustained neural resources in regions critical for WM. Mid-frontal theta is closely linked to top–down control, performance monitoring, selective attention, and executive functions that orchestrate maintenance and manipulation (Bush et al., 2000; Griesmayr et al., 2010; Hsieh & Ranganath, 2014; Mitchell et al., 2008; Onton et al., 2005; Wang et al., 2005). Age-related volumetric and functional declines in these systems likely constrain theta-based recruitment in older adults (Park & Reuter-Lorenz, 2009; Raz & Rodrigue, 2006; Tóth et al., 2014). Temporal theta contributions may additionally reflect sensory processing and hippocampal–cortical interactions underpinning maintenance and consolidation (Akiyama et al., 2017; Cabeza & Nyberg, 2000; Kamiński et al., 2017; Olson et al., 2006; Persson et al., 2012; Peterson & Thaut, 2002). In visuospatial contexts, temporal involvement plausibly supports spatial encoding and cognitive map formation—abilities that are especially vulnerable to ageing (Jeneson & Squire, 2012; Johnson et al., 2018; Kawasaki et al., 2014; Parvez & Khan, 2024).

Crucially, correlation analyses corroborated the functional relevance of theta modulation. In younger adults, increases in frontal, parietal, and temporal theta across the full trial predicted better performance in the Letter Span manipulation condition, and left temporal theta increases from delay to recall predicted accuracy in the Corsi retention condition. The absence of analogous correlations in older adults suggests that theta fluctuations indexed core WM mechanisms more directly in youth, whereas older adults may have relied on alternative, compensatory operations that are less tightly coupled to canonical theta networks (Cabeza et al., 2018; Pavlov & Kotchoubey, 2017, 2020; Reuter-Lorenz & Cappell, 2008). Theta dynamics also reflected load- and capacity-sensitive trajectories. Older adults consistently showed decreases from encoding to delay, compatible with limited capacity to sustain theta-mediated control as cognitive demands accumulate (Festini et al., 2018; Reuter-Lorenz & Cappell, 2008). Younger adults exhibited similar decreases only in the Corsi Test, alongside plateau effects in Letter Span manipulation, indicating proximity to saturation only under the more challenging visuospatial conditions. Notably, older adults displayed significant theta increases in the Letter Span but not in the Corsi Test, mirroring behavioural evidence that visuospatial load imposed disproportionate difficulty. These results align with the CRUNCH framework, suggesting that both groups recruit additional resources as demands rise (evidenced by delay-to-recall increases), but that older adults reach saturation at lower loads (reflected in encoding-to-delay decreases), especially in visuospatial contexts (Brookes et al., 2011; Heinrichs-Graham & Wilson, 2015; Kwon et al., 2015; Proskovec, Heinrichs-Graham, et al., 2019; Springer et al., 2023).

Importantly, despite robust age-group differences in theta, behavioural divergences appeared only in the Corsi Test. This dissociation indicates that theta alone does not account for preserved verbal WM in older adults; rather, other mechanisms likely supported performance, particularly in the Letter Span, where older adults matched younger adults despite diminished theta recruitment.

#### Alpha Power Changes

Alpha oscillations offered a complementary perspective, revealing task- and phase-dependent compensatory dynamics, particularly in older adults. In the Letter Span, older adults showed significantly stronger lower and upper alpha suppression than younger adults from delay to recall and across the full trial in both conditions. This pattern is consistent with reports of age-related alpha decreases during WM, often interpreted as increased cortical excitability and resource allocation in the service of maintenance, updating, and retrieval under higher demands (Pesonen et al., 2007; Proskovec et al., 2019a, 2019b; Scharinger et al., 2017; Sghirripa et al., 2021; Springer et al., 2023). Notably, despite the younger group’s theta advantages, the absence of a behavioural difference in the Letter Span suggests that enhanced alpha suppression in older adults played a compensatory role, enabling performance to be sustained (Angel et al., 2016; Klimesch, 1997; Klimesch et al., 1996; Michels et al., 2010; Murphy et al., 2020; Roux & Uhlhaas, 2014; van Ede, 2018).

Correlation analyses support this interpretation: in older adults, stronger lower alpha decreases over right parietal regions from encoding to delay predicted superior performance in the manipulation condition. Although right parietal cortex is not canonical for verbal processing, its increased contribution is consistent with compensatory redistribution and the HAROLD framework, wherein older adults recruit additional, often contralateral, regions to offset declining efficiency in typical networks (Cabeza et al., 2002, 2018; Klimesch, 1999; Klimesch et al., 2007; Pavlov & Kotchoubey, 2021; Scharinger et al., 2017; Stipacek et al., 2003).

In the Corsi Test, a different picture emerged. During retention, younger adults showed stronger upper alpha decreases in right frontal, temporal, and parieto–occipital regions, consistent with efficient, lateralised visuospatial recruitment and mirroring their behavioural advantage (Bonnefond & Jensen, 2012; Manza et al., 2014; Meltzer et al., 2008; Proskovec et al., 2019b; Wianda & Ross, 2019). Conversely, during manipulation, older adults exhibited stronger mid-frontal upper alpha suppression from delay to recall, yet this was accompanied by poorer performance—suggesting that the suppression indexed compensatory effort rather than efficient processing (Hsu & Hämäläinen, 2022; Kardan et al., 2020; Oberauer et al., 2018).

Additionally, both age groups displayed significant increases in posterior upper alpha from encoding to delay in the Corsi Test, contrasting with the suppression pattern observed in the Letter Span. This divergence implies a strategic “closed-gate” stance to protect stored visuospatial representations against interference, rather than prioritising continuous updating (Bonnefond & Jensen, 2012; Heinrichs-Graham & Wilson, 2015; Jensen et al., 2002; Jiang et al., 2015; Michels et al., 2008; Zhozhikashvili et al., 2022). In older adults, stronger right posterior alpha enhancement predicted better retention performance, underscoring a protective role for posterior alpha in safeguarding visuospatial information under sustained load (Bonnefond & Jensen, 2012; Manza et al., 2014; Meltzer et al., 2008; Proskovec et al., 2019b; Wianda & Ross, 2019).

Functionally, these task-contingent alpha patterns point to distinct strategies. In the Letter Span, sustained alpha suppression during delay is consistent with an “open gate” facilitating rehearsal and updating (D’Ardenne et al., 2012; Hazy et al., 2006; Katahira et al., 2018; Manza et al., 2014), and with continued engagement of the phonological loop over extended delays (7.5–10 s), even in retention trials without explicit updating demands (Awh et al., 1996; Baddeley & Hitch, 2019; Friederici et al., 2003; Paulesu et al., 1993; Riley & Constantinidis, 2016; Vallar et al., 1997). By contrast, in the Corsi Test, alpha increases indicate a “closed gate” strategy prioritising the protection of stored visuospatial content, consistent with higher difficulty and early emergence of capacity pressures (D’Ardenne et al.,. This interpretation converges 2012; Hazy et al., 2006; Katahira et al., 2018; Manza et al., 2014) with the theta saturation observed from encoding to delay in both groups, implying that task difficulty began to constrain resources early in the trial.

Notably, while group differences in alpha power emerged most clearly during the recall phase across both tasks, significant correlations between neural activity and performance were evident only during the delay period. This pattern suggests that older adults may have recruited compensatory mechanisms during maintenance and manipulation, despite the most pronounced alpha modulations occurring later at retrieval (Clements et al., 2023; Edin et al., 2009; Gajewski & Falkenstein, 2014; Gazzaley & Nobre, 2012; Kim, 2019).

Finally, evidence for functional differentiation across alpha sub-bands was apparent. In the Letter Span, lower alpha suppression aligned with rehearsal and updating processes, whereas upper alpha suppression may additionally index semantic activation supporting letter–word associations (Acheson & MacDonald, 2009; Klimesch, 1997; Klimesch et al., 2005, 2006). In the Corsi Test, selective modulation of upper alpha—rather than lower alpha—favoured active maintenance over global inhibition, consistent with the absence of external distractors and the need to stabilise internal representations over long delays and high load (Bonnefond & Jensen, 2012; Jensen & Mazaheri, 2010; Palva & Palva, 2007; Riddle et al., 2020; Sauseng et al., 2005; Scheeringa et al., 2009; Tuladhar et al., 2007).

In sum, older adults consistently exhibited stronger alpha suppression, but its functional significance varied by task: compensatory and performance-sustaining in the Letter Span, less effective under the heavier visuospatial burden of the Corsi Test. Alpha thus provided a flexible mechanism that complemented theta: whereas theta predicted efficiency and performance chiefly in younger adults, alpha modulation served as a principal compensatory channel in older adults.

### Limitations, Strengths, and Future Directions

Several limitations should be acknowledged. First, although EEG offers excellent temporal resolution, its spatial precision is limited relative to MEG, and cluster-based permutation analyses constrain anatomical localisation (Heinrichs-Graham & Wilson, 2015; Leenders et al., 2018; Proskovec et al., 2016; Proskovec, Heinrichs-Graham, et al., 2019; Proskovec, Wiesman, et al., 2019; Springer et al., 2023). Second, alternative approaches—source reconstruction, ERS/ERD quantification, or connectivity analyses—might yield complementary insights, but were constrained by the number of trials permitted by standardised WM test scoring. Third, the sample size limited modelling of inter-individual differences; prior work suggests substantial variability between high- and low-performers that could shape oscillatory–behavioural coupling.

Nonetheless, the study possesses notable strengths. Frequency bands were tailored to each participant’s individual alpha peak for each WM phase, accommodating inter-individual and age-related variability. The task architecture delineated encoding, delay (retention/manipulation), and recall, enabling phase-specific inferences. The inclusion of verbal and visuospatial domains—each crossed with retention and manipulation—permitted evaluation of compensation across modalities and operations. Finally, assessing memory performance along a continuum (rather than dichotomously) increased sensitivity to gradations in storage and manipulation capacity.

Future work should couple EEG with source modelling to refine spatial inferences, increase trial numbers to support richer time–frequency and connectivity analyses, and recruit larger cohorts to parse individual differences and performance strata (Tagliabue & Mazza, 2021). Equating task difficulty across age groups would clarify whether observed differences reflect capacity per se or strategic choices under asymmetric load. Distinguishing lower versus upper alpha effects more systematically could further elucidate their distinct contributions to rehearsal, protection, and interference control. Finally, administering the Letter Span and Corsi tasks in separate sessions may mitigate fatigue or motivational drifts, thereby improving data quality.

## CONCLUSION

This study advances understanding of age-related WM by integrating behavioural performance with phase-specific oscillatory dynamics across verbal and visuospatial tasks, particularly in maintenance/manipulation and recall phases. Older adults maintained performance in the Letter Span but showed clear deficits in the Corsi Test, indicating domain specific vulnerability in visuospatial WM. Task complexity exerted strong, group-general effects, underscoring that manipulation demands constrain performance irrespective of age.

At the neural level, theta and alpha revealed complementary roles. Younger adults exhibited stronger mid-frontal and temporal theta power that predicted performance in high-demand contexts, consistent with efficient engagement of core WM mechanisms. Older adults showed reduced theta—suggestive of earlier saturation—alongside enhanced alpha suppression and task-contingent alpha increases, consistent with compensatory resource allocation and protection of stored representations, particularly during the delay period (maintenance / manipulation).

Collectively, these findings refine cognitive ageing models by demonstrating that oscillatory dynamics mediate a dynamic balance between efficiency and compensation that is both task-specific and phase-dependent. Theta primarily supports efficient processing in youth, whereas alpha provides a flexible compensatory scaffold that can sustain performance when efficiency wanes. Identifying when and where these mechanisms operate offers mechanistic targets for interventions—such as non-invasive brain stimulation or cognitive training—aimed at bolstering WM in ageing populations.

## SUPPLEMENTARY MATERIAL

### Additional information about training and near-transfer WM tasks

#### 1. OSPAN

⍰ Arithmetic problems included only sums, subtractions, multiplications and divisions, and they were presented with their proposed solution (e.g., (5 x 7) + 3 = 37).
⍰ During recall, participants had no time limit, but they were not allowed to correct typing mistakes. This ensured that participants typed the digits in the correct order rather than entering them randomly to delay memory decay.
⍰ Before training, all participants practised each task component separately and then together. This warm-up consisted of:
  ∘ Two trials of digit recall only (two digits without arithmetic problems).
  ∘ Two trials of only arithmetic problems (no time limit) o Four trials of the combined task.
    - The warm-up was repeated at the start of each training session, and training began only once participants demonstrated familiarity with both components.
⍰ Each training session consisted of three blocks of ten trials each. All participants started with a sequence of two digits.
  - During the task, the number of digits increased or decreased based on performance:
    ∘ If a participant recalled the digits correctly twice in a row at the same level, another digit was added to the sequence for at least the next two trials.
    ∘ If they answered incorrectly twice in a row at the same level, a digit was removed from the sequence for at least the next two trials.
    ∘ If they had mixed results (one correct and one incorrect, or vice versa), the sequence length stable until one of the other conditions was met.
⍰ Two metrics were extracted for each session: raw scores and the highest level completed (Conway et al., 2005). Raw scores were calculated by multiplying the number of correct answers by their respective level and summing across all blocks. The highest level completed was defined as the highest sequence length at which both series of digits were correctly recalled.

#### 2. N-back

⍰ Participants were explicitly instructed to respond on every trial and to rely on memory rather than guessing.
⍰ Before training, all participants completed four short practice blocks of 10 trials each to familiarise themselves with the task. The first two blocks were 1-back, and the last two were 2-back. After each practice block, participants received feedback on their accuracy, which reflected the percentage of correct answers. Task difficulty during training was adjusted according to this performance.
⍰ Each training session consisted of 14 blocks of 20 trials each, beginning at 1-back. 70% of the trials were non-targets (did not match location) and 30% were targets (matched location).
⍰ Difficulty was adapted according to the following rules
  ∘ If a participant scored 90% or higher, the level increased by one (e.g., from 1-back to 2-back)
  ∘ If they scored between 71% and 89%, they remained at the same level
  ∘ If the score was 70% or lower, they dropped one level (e.g., from 3-back to 2-back), except for 1-back, where they stayed at the same level.
⍰ These percentages ensured that participants relied on their memory skills and did not advance by chance.
⍰ Two metrics were extracted for each training session: the raw score and the highest level reached (e.g., 2-back, 3-back). Raw scores were calculated by multiplying the number of correct answers by the level of the respective block and summing across all blocks. The task took approximately 20 minutes to complete.

#### 3. WM tests: OSPAN and N-back

⍰ The set of 12 consonants (C, F, H, J, L, N, K, P, Q, R, V, W) was used in the Letter Span to minimise the possibility of word formation and to reduce rehearsal strategies.
⍰ The grid layout in both tasks remained constant across trials to avoid introducing additional visual search demands.
⍰ The zero key could be used once per trial in both tasks if a participant forgot an element, ensuring participants completed sequences even with partial recall.
⍰ Scoring was based on one point per correctly recalled element to maintain motivation and engagement, even when sequences were only partially remembered.
⍰ Before recruitment, both tests were piloted with a group of ten old adults and five young adults to adjust for difficulty factors such as the number of elements to recall, the duration of the phases, and the response method.
⍰ Stimuli were also tested. For instance, in the Letter Span, participants had difficulties differentiating the “P” from the “T” or the “F” from the “S” thus the “T” and the “S” were excluded.

### Additional information about EEG preprocessing

#### 1. Channel and Component Rejection Criteria

⍰ Channels rejected if flat >5 seconds, poorly correlated (&80%) with neighbours, or extreme amplitude (+/- 4 SD).
⍰ ICA components rejected if >80% eye or muscle activity.
⍰ Maximum 10% of channels and 20% of ICA components could be rejected to avoid excessive data loss.
⍰ Channel and component rejection criteria were based on previous pipelines (Gil Ávila et al., 2023; Pernet et al., 2021).

#### 2. Epoching Details

⍰ Encoding: 0 to 8.5 seconds relative to stimulus onset.
  - Delay – retention: 0 to 7.5 seconds.
  - Delay – manipulation: 0–10 seconds.
  - Recall – retention/manipulation: 0 to 4 seconds to standardise across participants due to variable response times.
  - Segmenting later into 1-s subsegments improves artifact detection and allows consistent spectral analysis across WM phases.

#### 3. Trial Inclusion/Exclusion

⍰ Minimum 50% accuracy per trial required.
⍰ Participants excluded from EEG analysis of specific conditions if &25% valid trials.
⍰ Participants excluded after this stage:
  - Letter Span – retention: 0 Young – 0 Old
  - Letter Span – manipulation: 6 Young – 2 Old
  - Corsi Test – retention: 1 Young – 4 Old
  - Corsi Test – manipulation: 1 Young – 11 Old
⍰ Minimum 15 valid subsegments per phase required for inclusion.
  - No participants excluded after this stage
⍰ One participant from each age group was excluded from all the tasks due to EEG recording errors.

### Additional information about statistical analysis

#### 1. EEG Analysis

⍰ Power spectra were averaged per participant (channel × frequency) and log-transformed.
  - Frequency bands were defined relative to individual alpha frequency (IAF; Klimesch, 1999):
    ∘ Theta: 5 Hz below IAF to 1 Hz below low-alpha lower bound o Lower alpha: IAF to 2 Hz below IAF
    ∘ Upper alpha: 2 Hz above IAF
  - Monte Carlo simulations were used to generate a null distribution of the cluster-sum statistic. Then, cluster-based permutation independent sample t-tests (10,000 iterations) were used to compare young vs old participants’ power changes across WM phases (encoding – delay, delay – recall, encoding – recall).
⍰ The highest sum of t-values across all clusters in each permutation were taken and added to a distribution of cluster statistics. Effects observed in the data were compared against this cluster distribution. A *p* value of 0.05 was considered as the threshold for significance.
⍰ A triangulation method was employed for clustering using the function *ft_prepare_neighbours* in Fieldtrip with a minimum of at least three channels per cluster.
⍰ Relative power (phase differences) was calculated by subtracting the power of one WM phase from the other.
⍰ Six relative phase comparisons: encoding vs delay-retention, delay-retention vs recall retention, encoding vs recall-retention, encoding vs delay-manipulation, delay-manipulation vs recall-manipulation, encoding vs recall-manipulation.
⍰ Effect sizes: upper and lower bounds of Cohen’s *d* per cluster (Meyer et al., 2021).

#### 2. Neural–behaviour correlations

- Permutation-based Spearman correlations (Python 3.10) were conducted using electrode clusters.
- EEG variables were first grouped by frequency band, WM phase, and electrode, then clustered spatially:
  ∘ Left frontal: F1, F3, F5, FC3
  ∘ Right frontal: F2, F4, F6, FC4
  ∘ Mid-frontal: Fz, Cz, FCz
  ∘ Left parietal: P1, P3, P5, CP3, PO3
  ∘ Right parietal: P2, P4, P6, CP4, PO4
  ∘ Occipital: O1, O2, Oz
  ∘ Left temporal: T7, TP7, FT7, FT9
  ∘ Right temporal: T8, TP8, FT8, FT10
- ⍰ FDR correction for multiple comparisons; bootstrapped 95% confidence intervals for robustness.

**Table S1.**
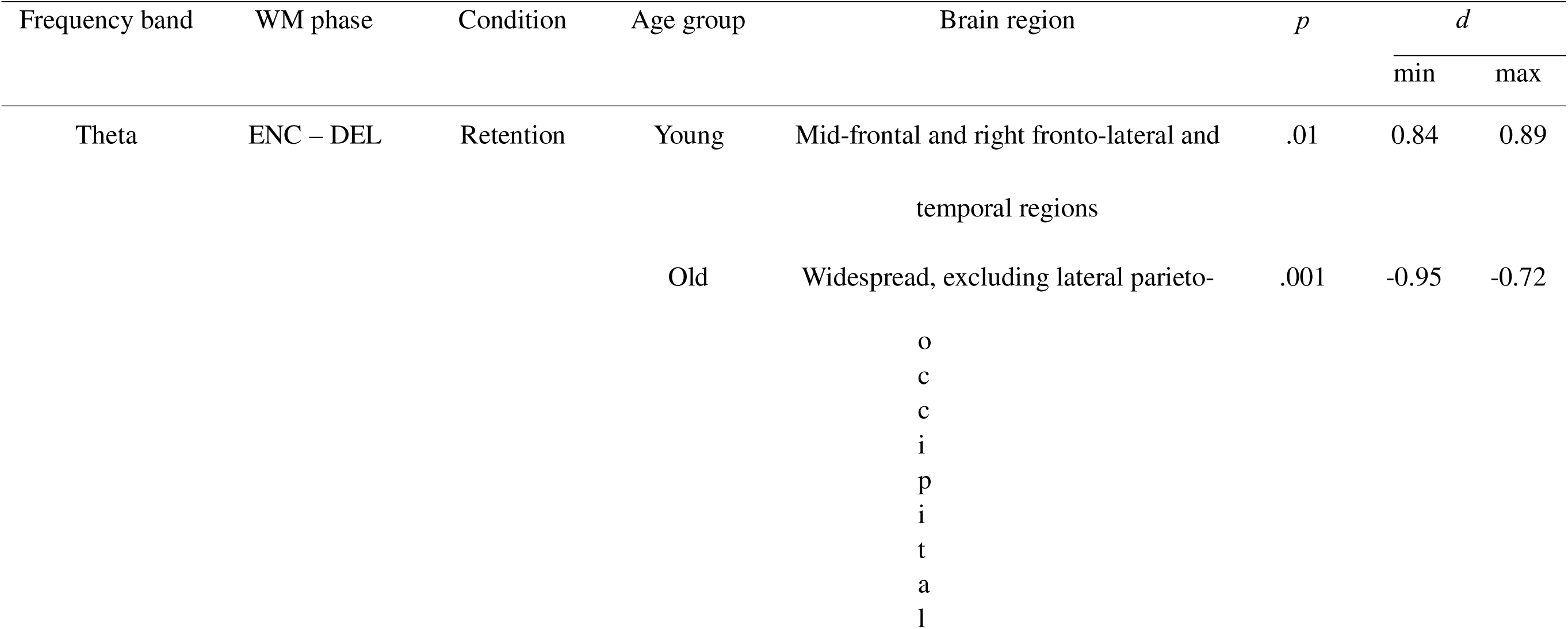
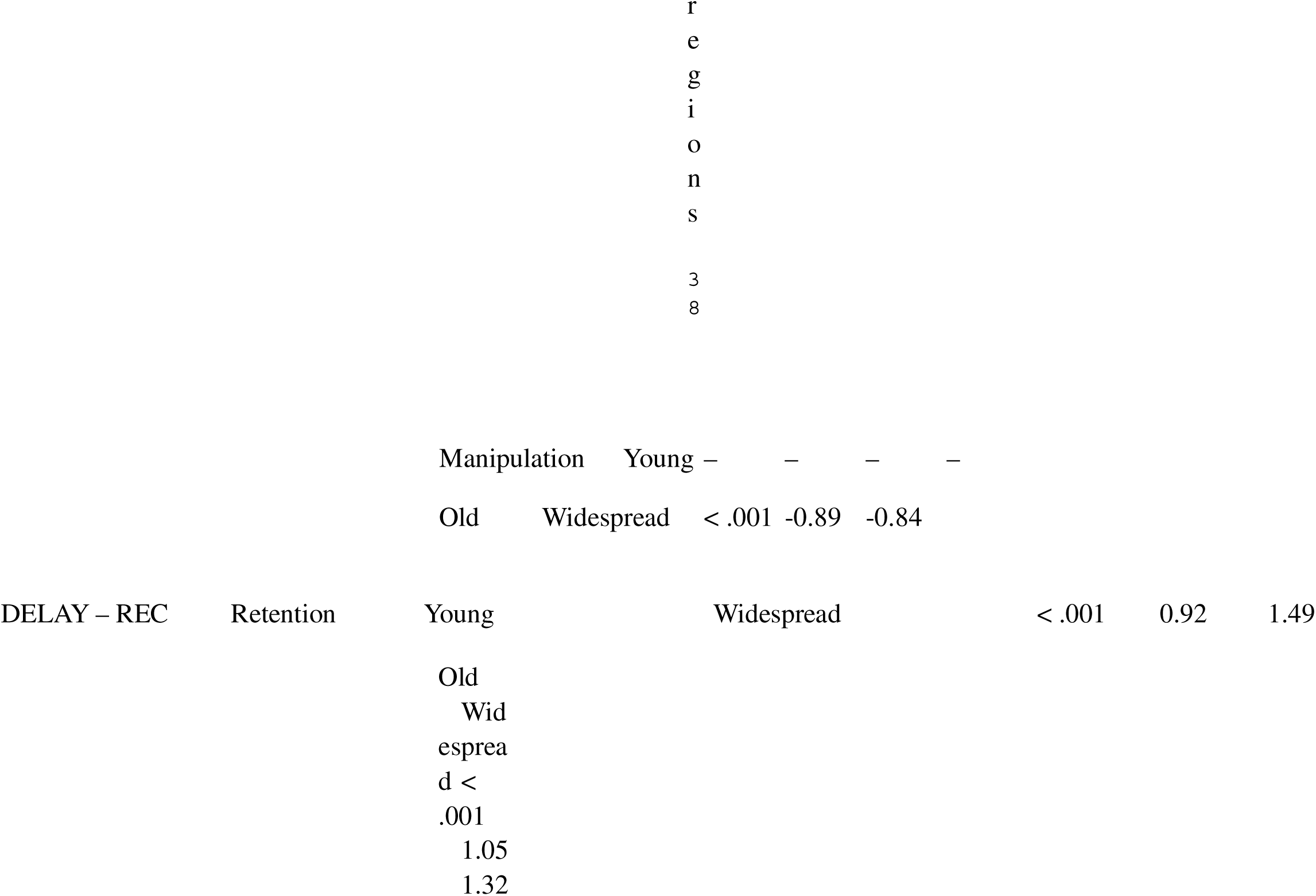
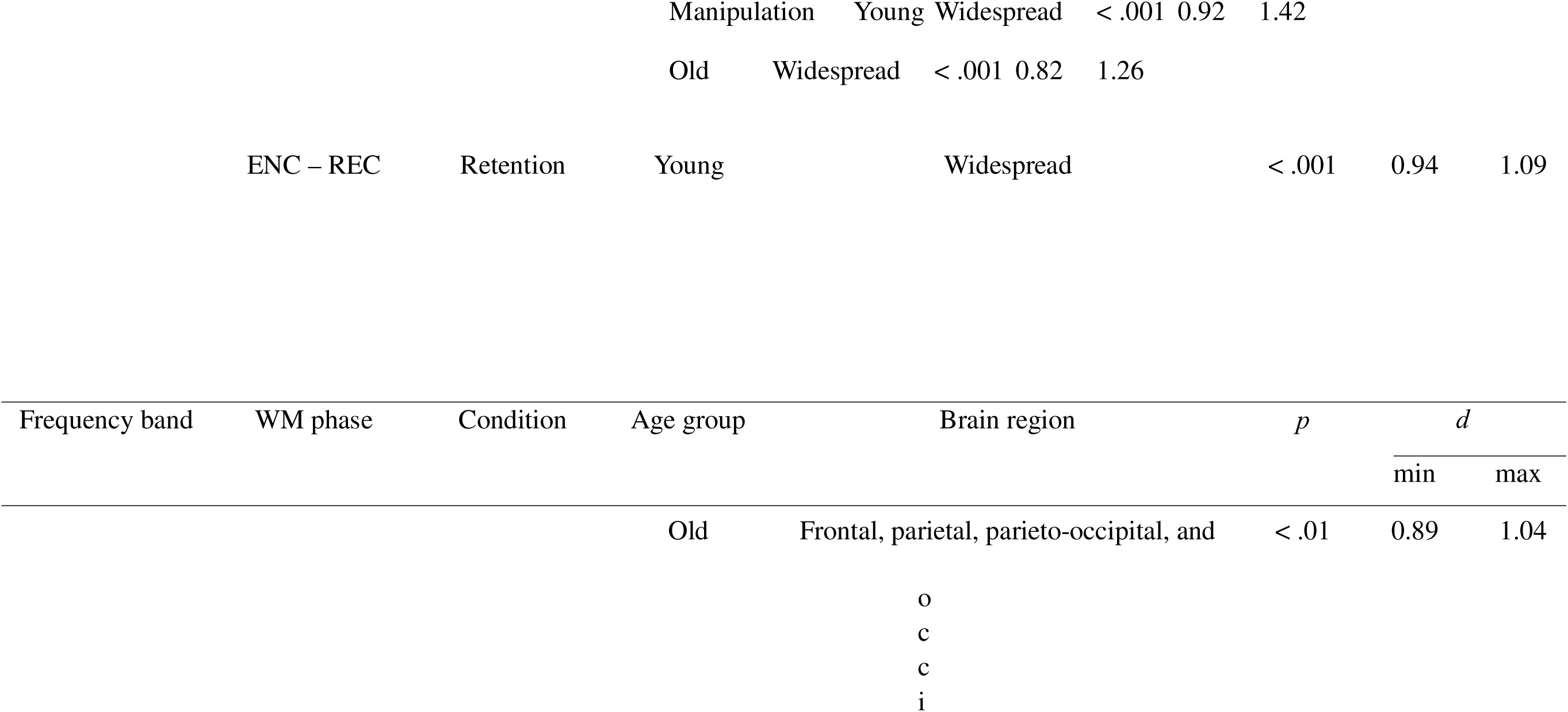
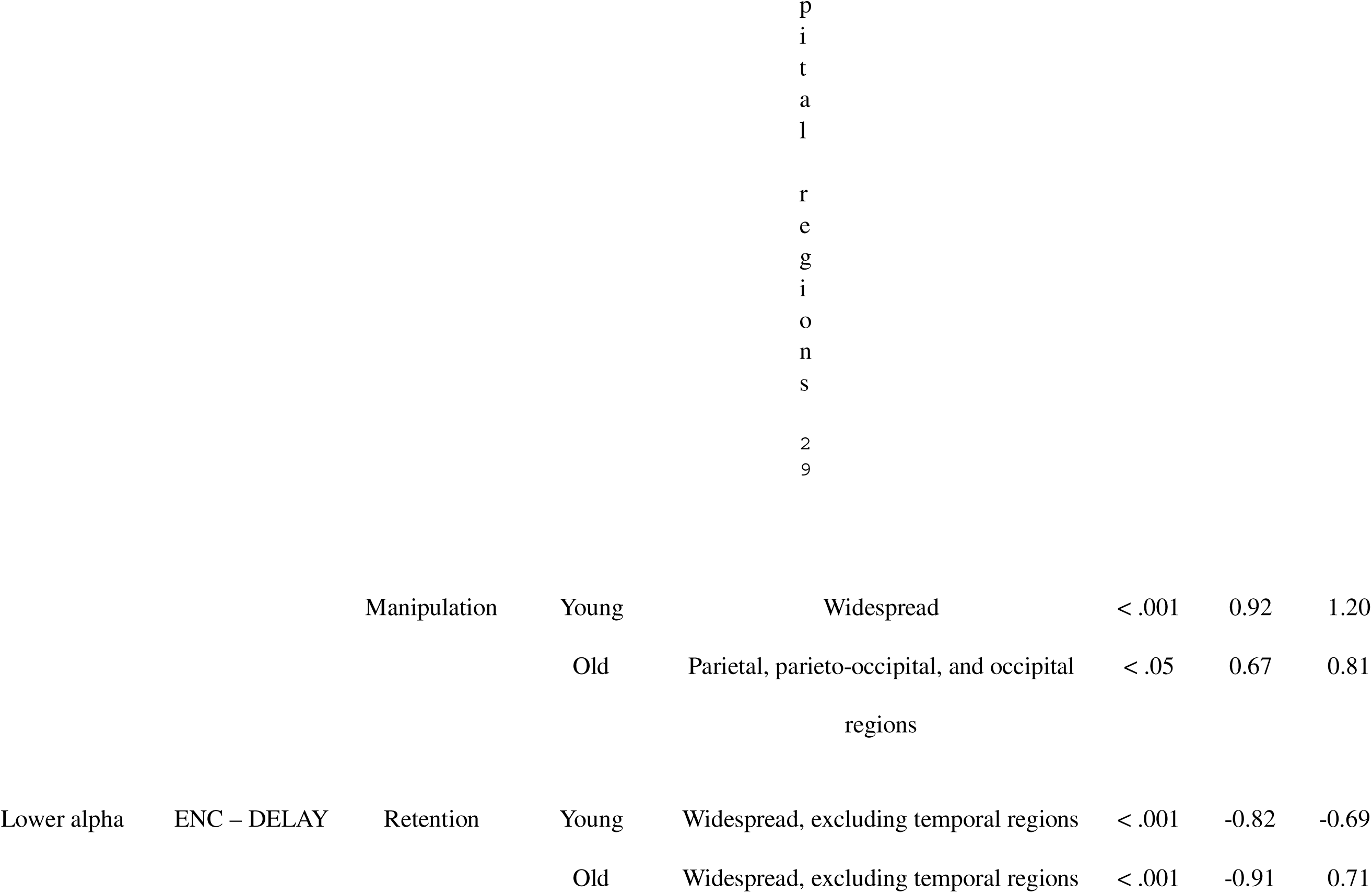
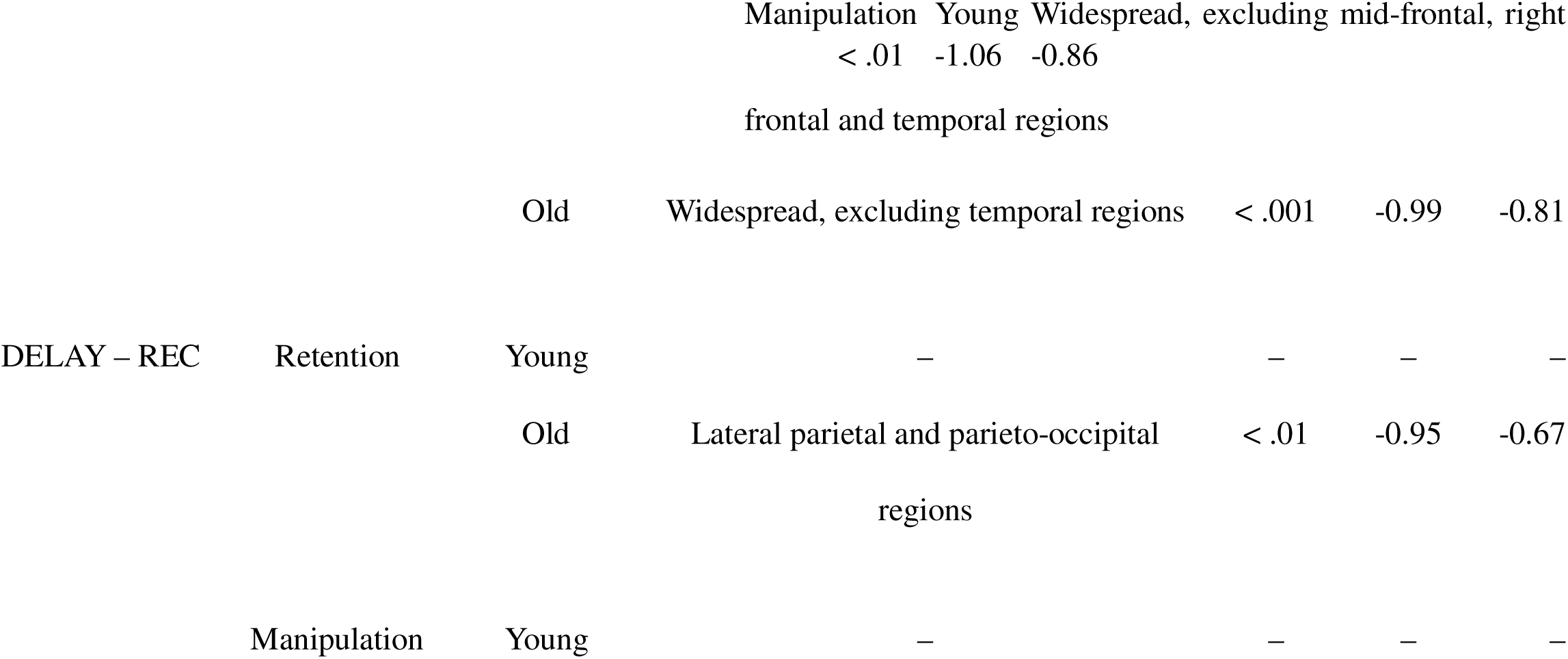
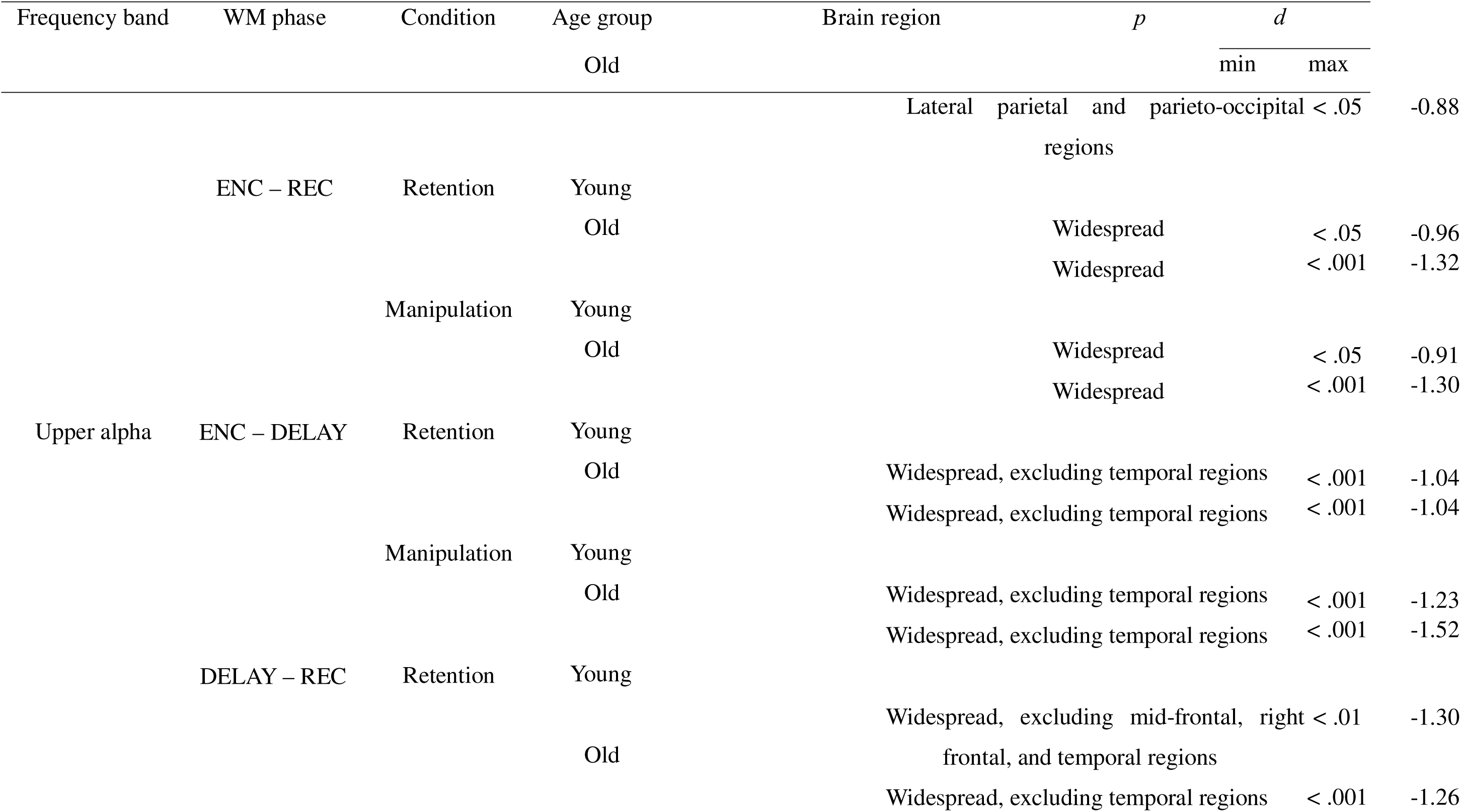
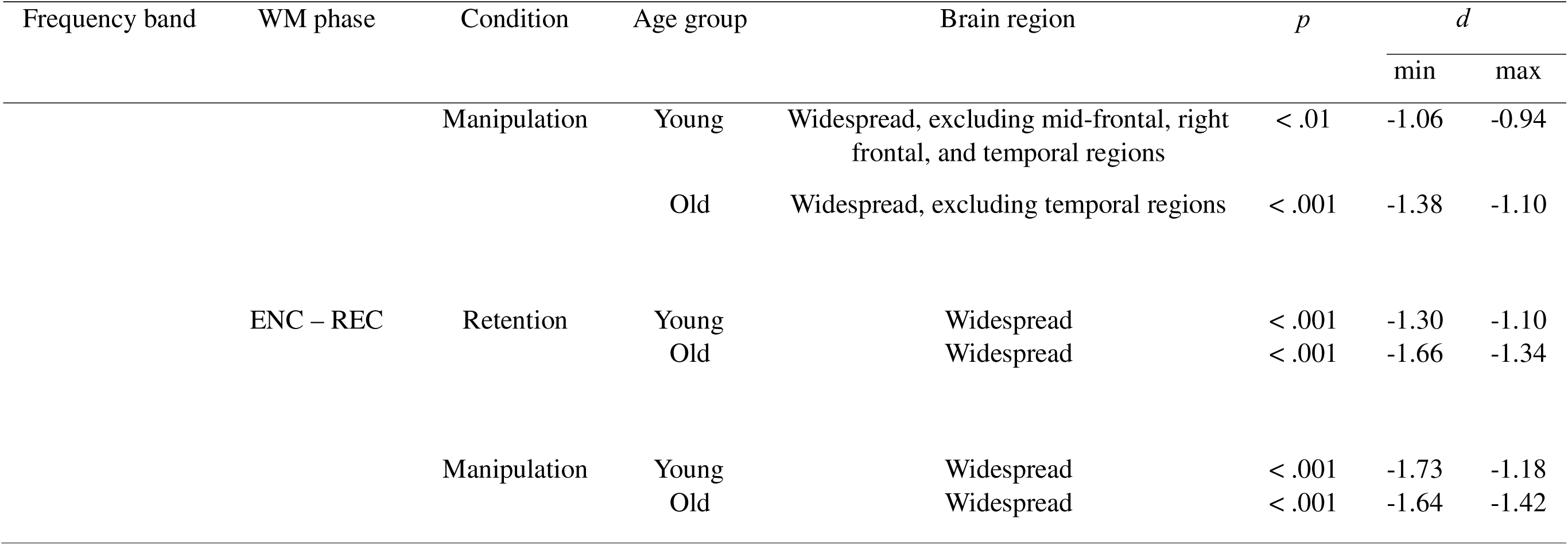
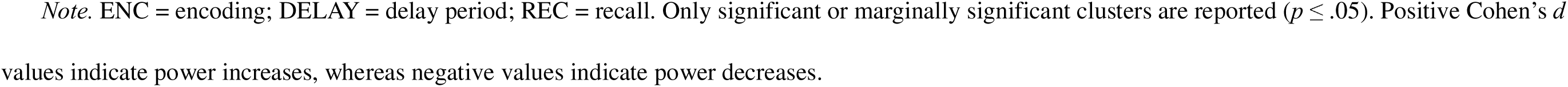
Exploration of oscillatory trends: significant differences in power changes within each age group across WM phases in the Letter Span.

**Table S2.**
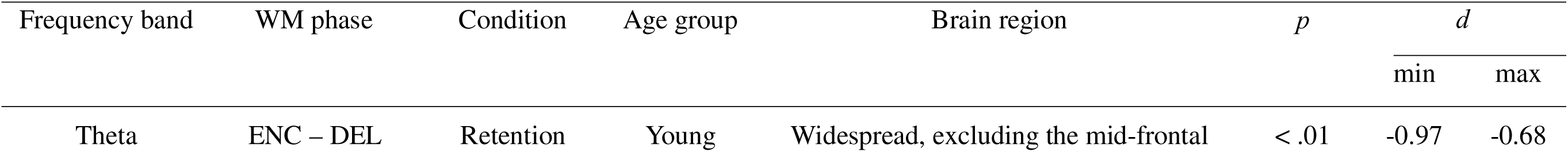
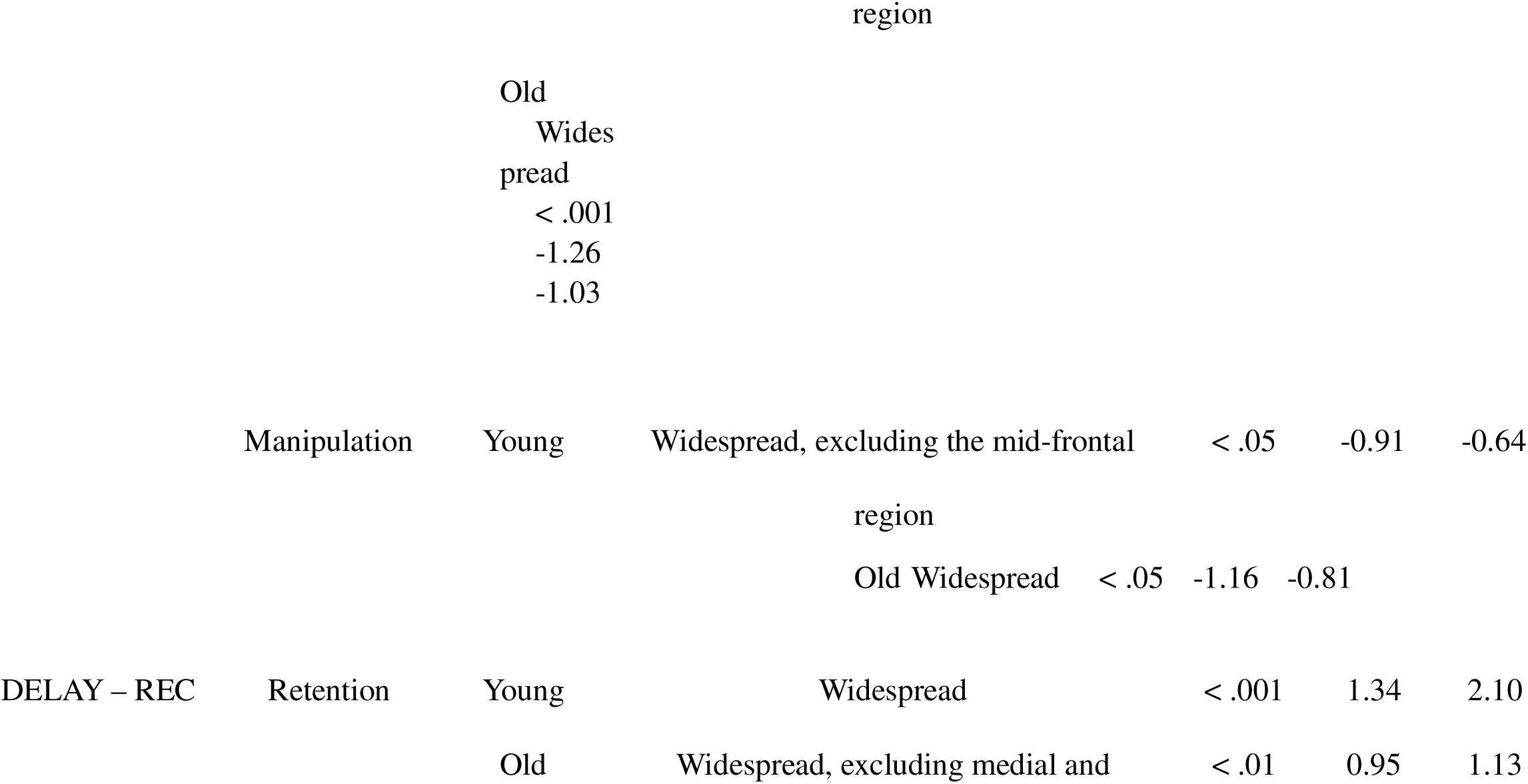
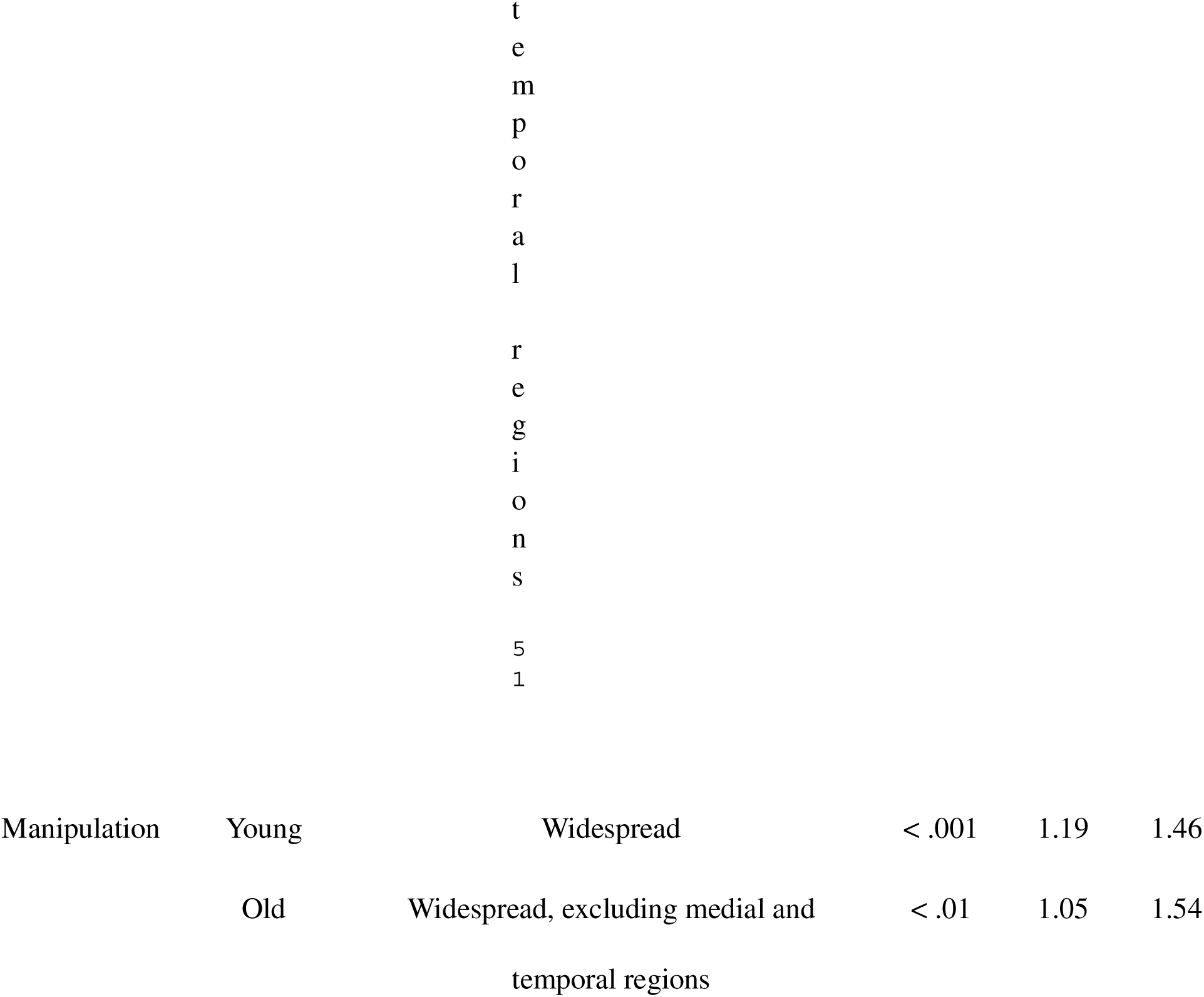
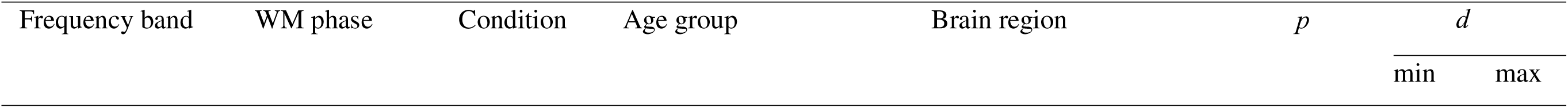
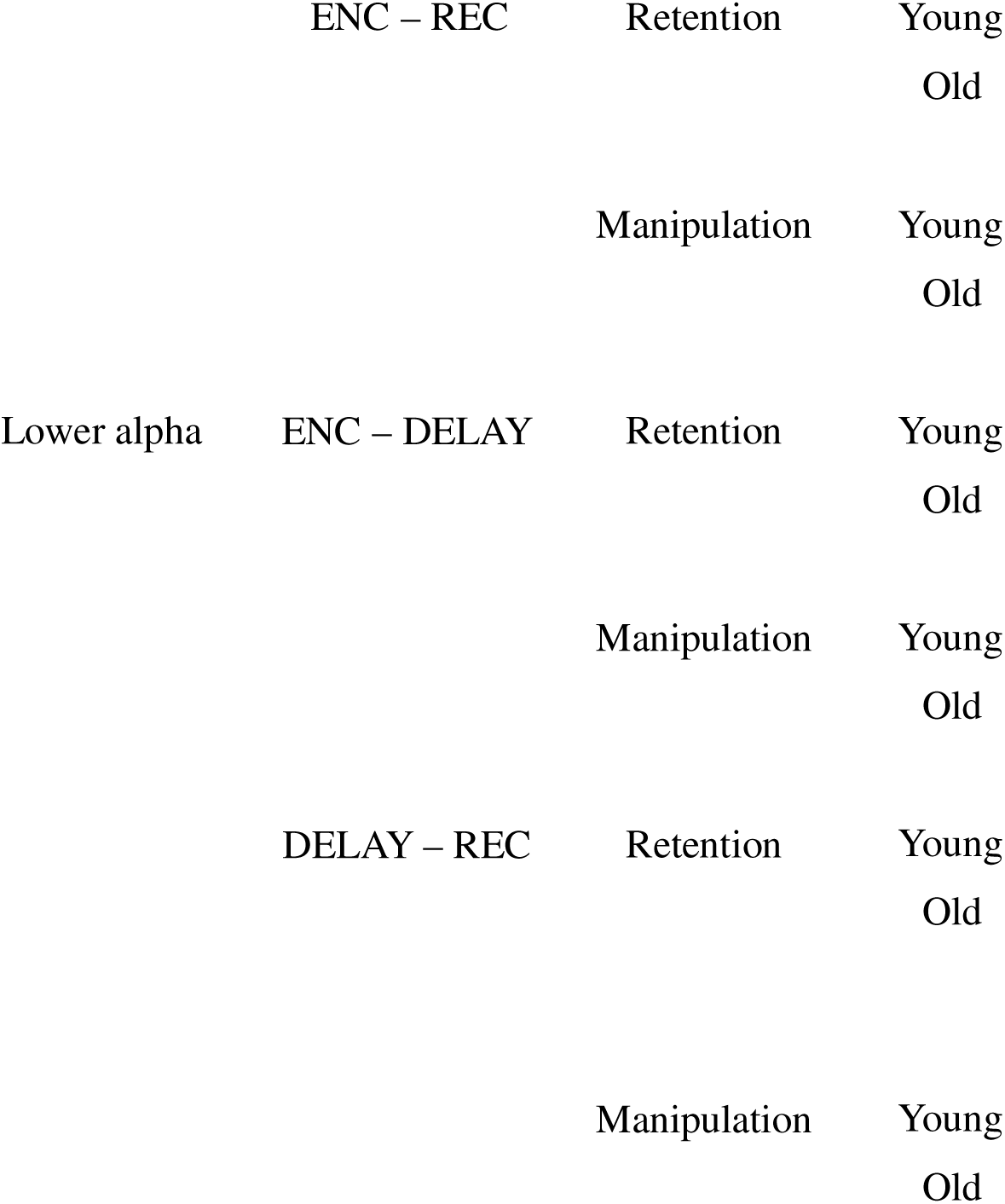
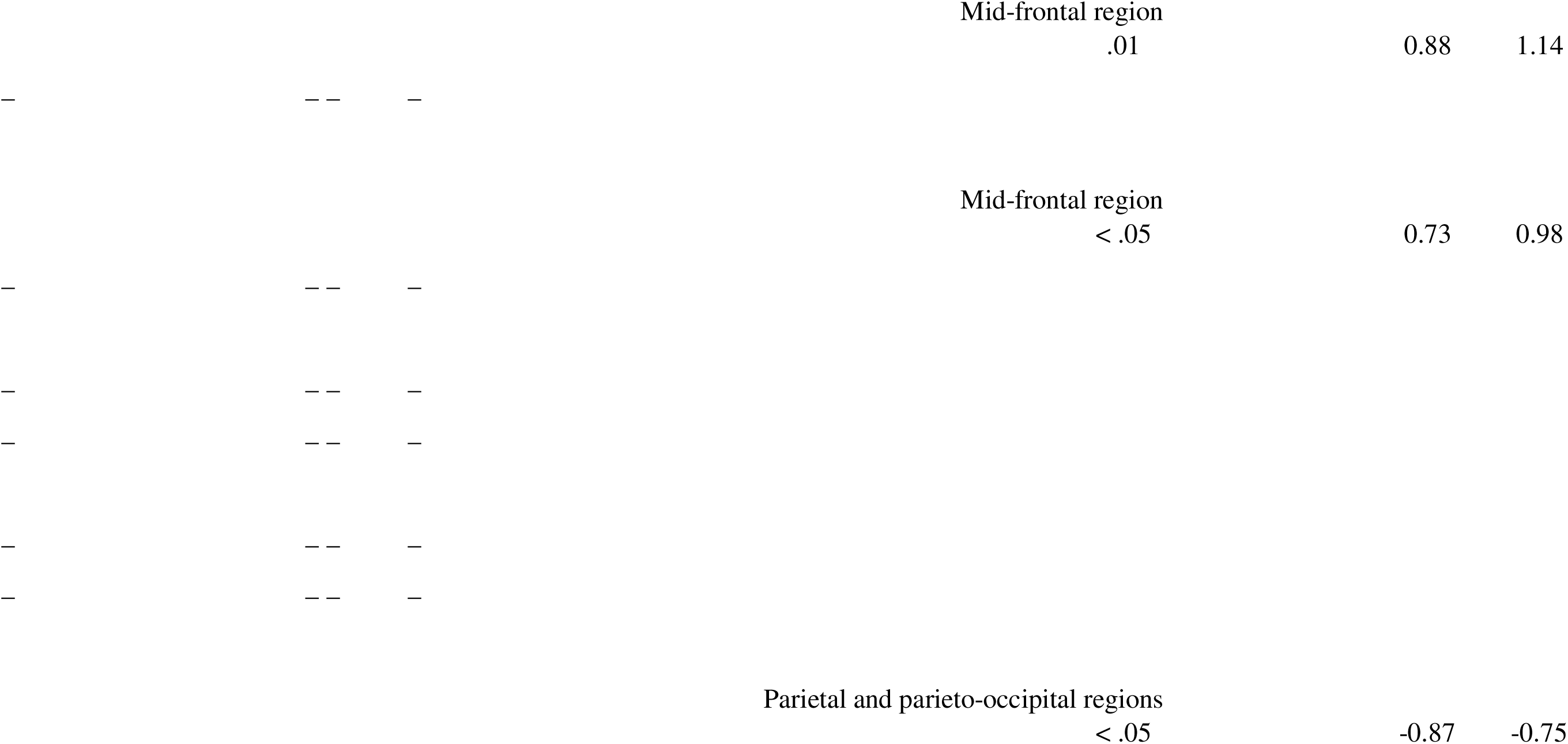
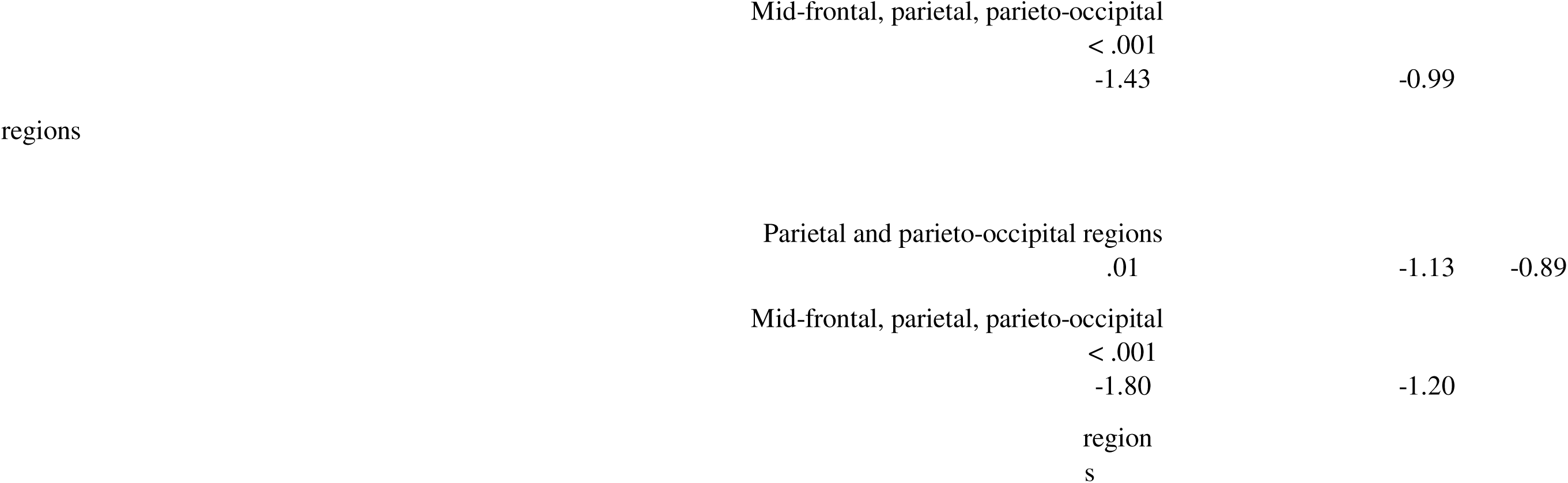
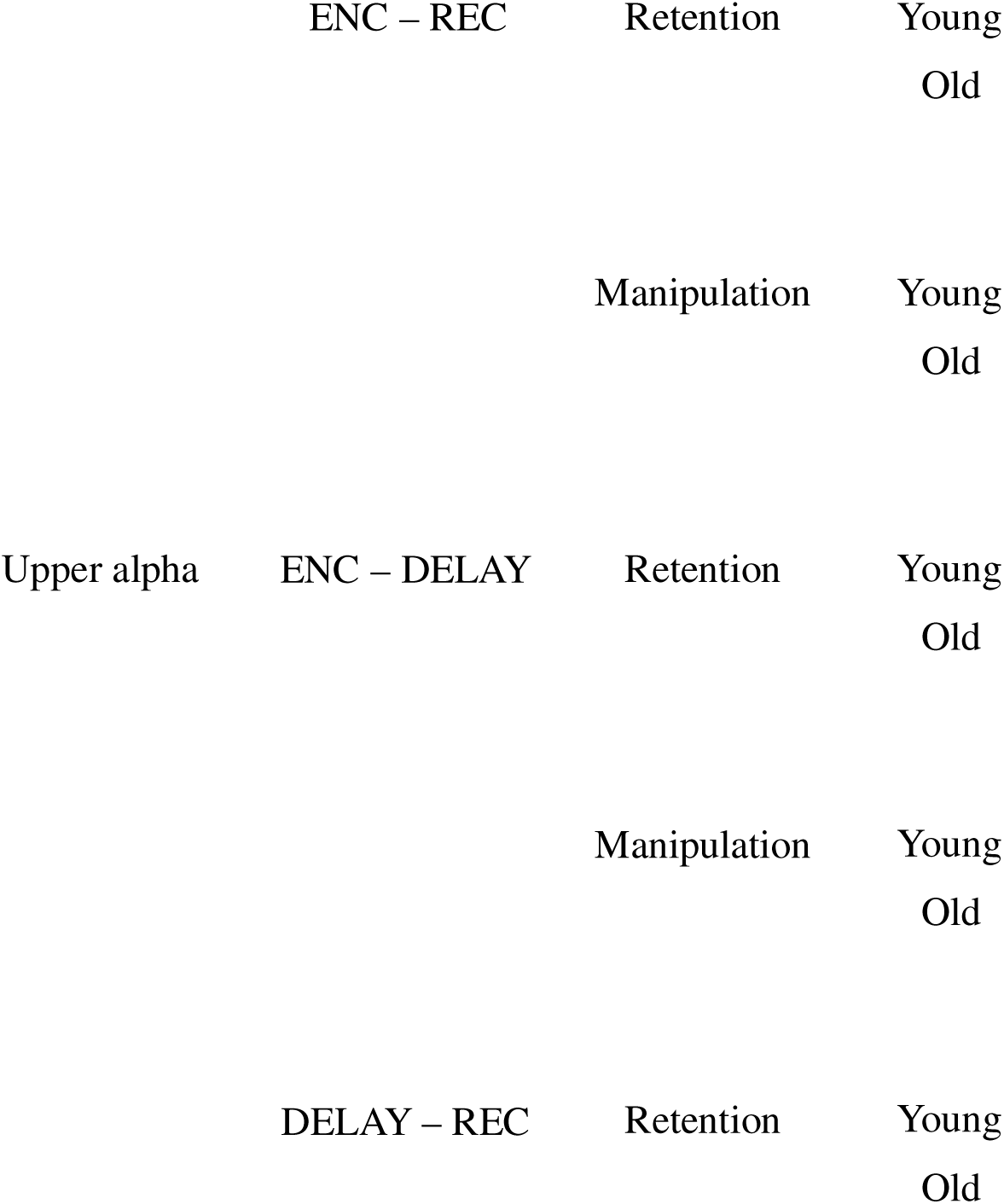
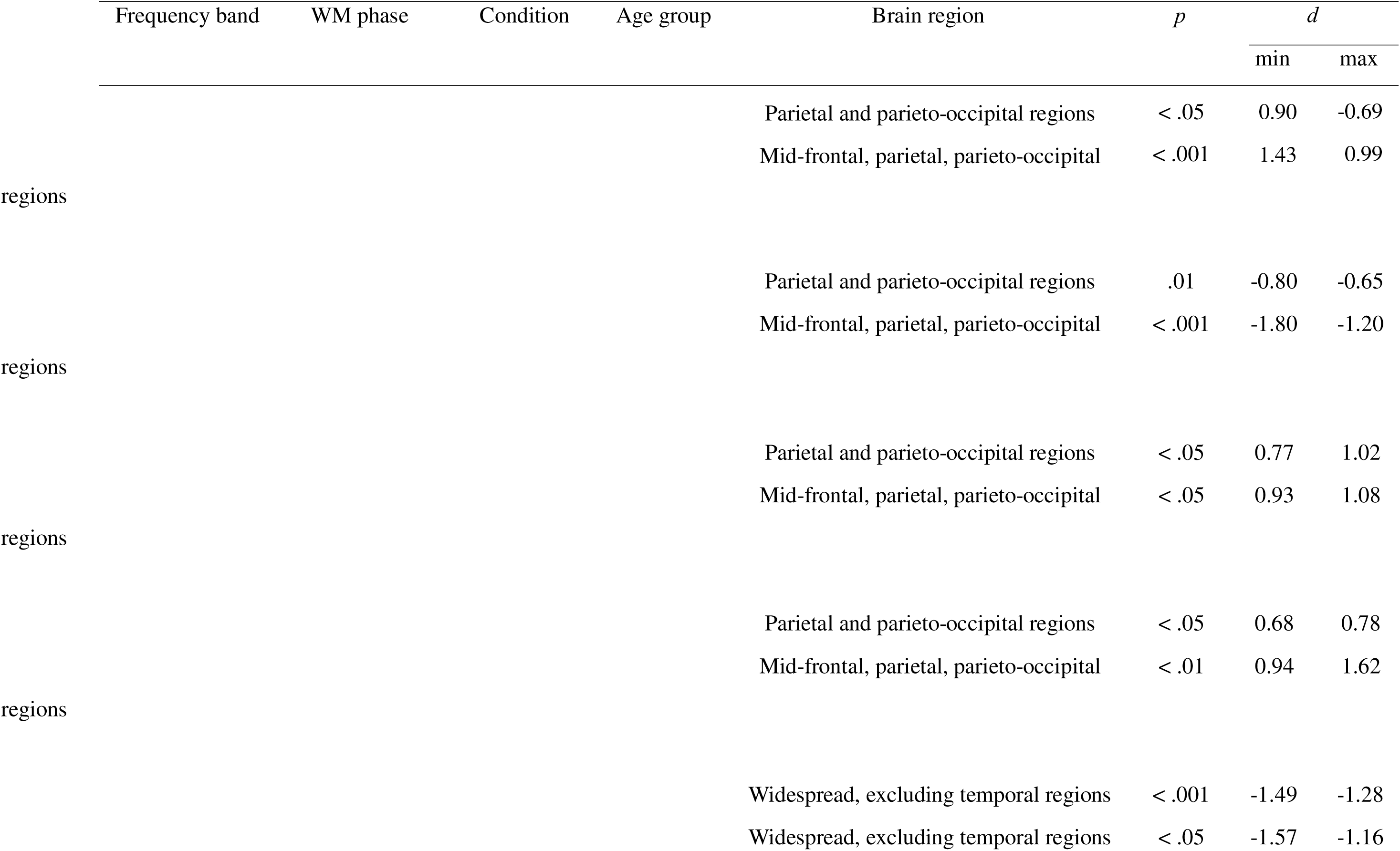
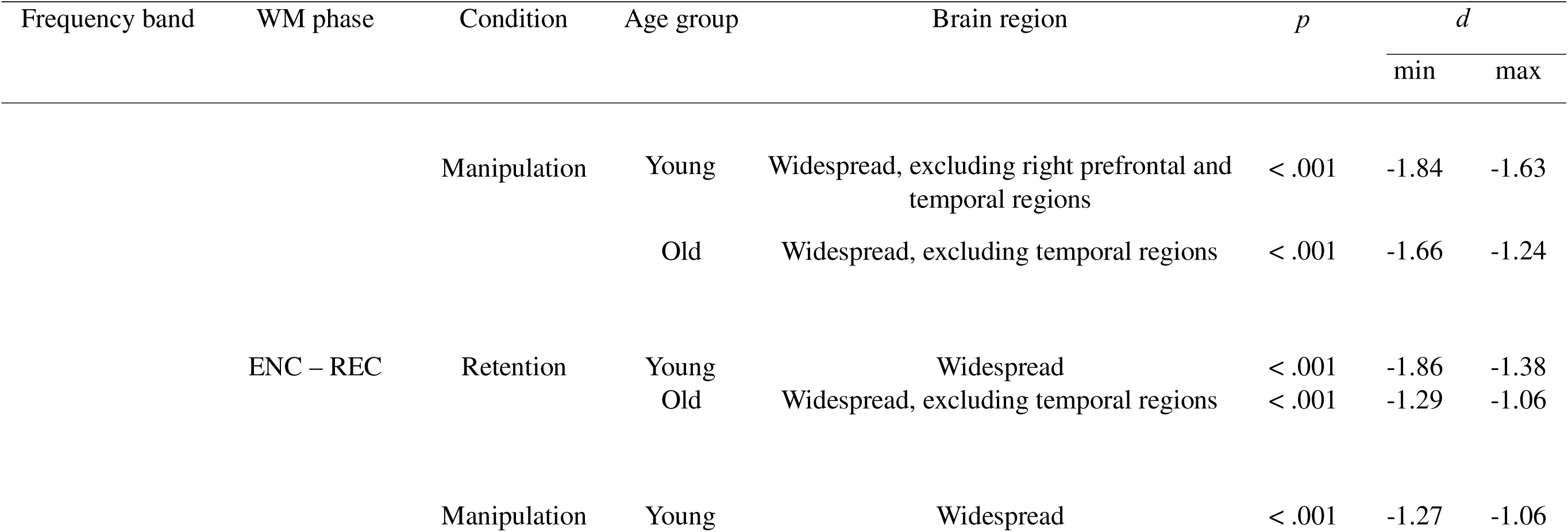
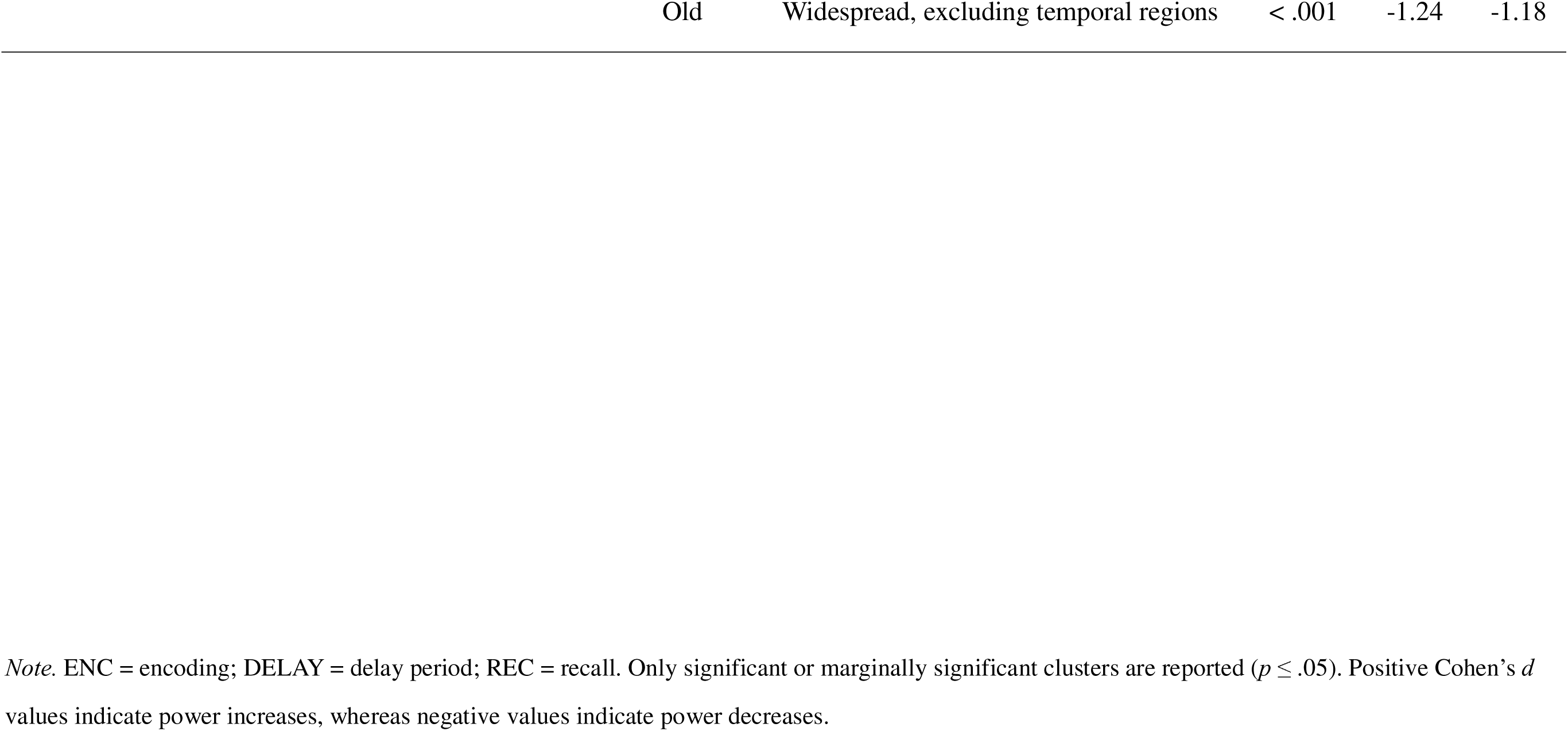
Exploration of oscillatory trends: significant differences in power changes within each age group across WM phases in the Corsi Test.

